# Global analysis of the cold-shock response in the model antibiotic producing actinomycete, *Streptomyces coelicolor* A3(2)

**DOI:** 10.1101/2025.04.11.648403

**Authors:** R. Tony Evans, Yongjae Lee, Andrew Hesketh, Byung-Kwan Cho, Colin P. Smith, Giselda Bucca

**Affiliations:** School of Applied Sciences, University of Brighton, Lewes Road, Brighton, UK; Graduate School of Engineering Biology, Korea Advanced Institute of Science and Technology, Daejeon 34141, Republic of Korea; KI for the Bioinnovation, Korea Advanced Institute of Science and Technology, Daejeon 34141, Republic of Korea; School of Immunology & Microbial Sciences, Faculty of Life Sciences & Medicine, Guy’s Hospital, Borough Wing, Great Maze Pond, King’s College London, London, SE1 9RT, UK; Present address: School of Biosciences, Faculty of Health and Medical Sciences, University of Surrey, Surrey, GU2 7XH, UK

**Keywords:** Translational control, polysome profiling, transcriptome, translatome, cold-shock response, *Streptomyces*, gene regulation, synthetic biology, iModulon

## Abstract

Cold-shock adaptation is essential for the survival of soil-dwelling sessile streptomycetes exposed to fluctuating environmental temperatures, yet the precise regulatory mechanisms underlying this response remain poorly understood. Here, we investigated the global transcriptional and translational responses of the model actinomycete, *Streptomyces coelicolor* A3(2), to cold-shock using integrated RNA-seq and polysome profiling. Cold-shock treatment in minimal liquid medium triggered significant transcriptional changes in 811 genes. Notably, three operons, encoding a CspA homologue, a DEAD-box helicase, and a cystathionine-β-synthase (CBS) domain-containing protein and/or a protein of unknown function (*SCO5921–SCO5918*, *SCO4684-SCO4686*, and *SCO3731-SCO3733*) were identified as central players of the cold-shock response, exhibiting up to 2,000-fold transcriptional induction. Systems-level transcriptomic analysis further revealed the cold-shock induced activation of pathways associated with gluconeogenesis, coenzyme A metabolism, phenylacetate degradation, lipid raft remodelling, and extracellular functions, pointing to extensive metabolic reprogramming coordinated with membrane adaptation during cold acclimation. Polysome profiling unveiled strong translational potentiation of operonic genes downstream from the promoter proximal *cspA* homologue genes, a process potentially mediated by RNA secondary structures that overlap ribosome binding sites (RBSs). The pronounced induction of DEAD-box RNA helicases and CspA RNA chaperones is presumed to reflect their critical requirement for resolving excessive nucleic acid secondary structures inherent to the high G+C content genome of *Streptomyces* (>73% G+C), including the RBS-masking stem-loops within their own operons. Together, this study provides a comprehensive, system-level understanding of cold-shock adaptation in *Streptomyces*, highlighting a multi-layered regulatory architecture that couples metabolic reprogramming with RNA structure-dependent translational control to mitigate thermal stress.

**IMPORTANCE:** This study characterizes the cold-shock response of the model actinomycete, *S. coelicolor* A3(2), for the first time at both the transcriptome and translatome levels. Combined with a machine-learning based iModulon framework, our findings provide critical insights on both metabolic adaptation and multi-layered regulatory mechanisms, including transcriptional networks and RNA structure-dependent translational control. Beyond advancing our fundamental understanding of cold acclimation in *Streptomyces*, we identified several putative *cis*-acting regulatory elements within the intergenic regions between the primary cold-shock genes (*SCO4684* and *SCO5921*) and their downstream DEAD-box helicase-encoding genes. These regulatory elements, coupled with the exceptionally robust transcriptional and translational induction of the core cold-shock operons, significantly expands the synthetic biology toolkit for *Streptomyces*. Ultimately, these molecular components hold substantial potential for exploitation in optimizing and manipulating cryptic antibiotic biosynthetic gene clusters within this bacterial genus of considerable industrial importance.

## INTRODUCTION

Streptomycetes are notable for producing diverse secondary metabolites, including antibiotics, and for their unusual developmental biology and high genomic GC content. Ubiquitous in soil, they inhabit highly dynamic and competitive ecosystems, requiring rapid responses to environmental fluctuations such as changes in pH, temperature, nutrient availability, or antibiotic exposure from competing organisms. We previously demonstrated that the streptomycete heat-shock response is largely post-transcriptionally controlled, enabling an accelerated physiological response to thermal stress (1). Among 74 more efficiently translated genes identified under heat-shock, the most enriched functional cluster comprised nucleic acid binding proteins, including six putative cold-shock proteins (CSPs): SCO0527, SCO3731, SCO3748, SCO4505, SCO4684 and SCO5921.

The cold-shock response in *Escherichia coli and Bacillus subtilis* has been extensively studied, revealing the central role of the cold-shock protein A (CspA) (2–5). However, no comparable systems-level studies have been reported for the model actinomycete, *Streptomyces coelicolor* A3(2). Post-transcriptional and translational regulation of *cspA* and other cold-shock mRNAs is well documented in *E. coli* (6). For instance, the *cspA* mRNA 5’-UTR functions as a thermosensor that stabilizes the transcript at low temperature, facilitating ribosome binding via a ‘downstream box’ sequence, while trans-acting factors such as IF3 and CspA itself, together with *cis*-acting UTR elements, further contribute to translational bias during cold acclimation (6–8). Additionally, the ribosome itself acts as a temperature sensor, adjusting its protein composition to yield a specialized translation apparatus optimized for cold-shock mRNA translation (2).

Mechanistically, Zhang et al showed a dramatic increase in global mRNA secondary structure within 30 min of cold-shock in *E. coli*, accompanied by a global arrest in protein translation due to reduced elongation rates and inhibited initiation (9). As cells acclimatize over 6 hours, structured RNA abundance decreases and translation gradually resumes. Two key proteins drive this acclimation: RNaseR, which degrades highly structured untranslated mRNAs, and CspA, an RNA chaperone that prevents inhibitory secondary structures and promotes translation initiation and elongation (9–11). More recently, rho-dependent transcription attenuation within the upstream regions of cold-shock genes was identified as an additional regulatory layer, which is relieved upon cold-shock and reactivated during acclimation as CspA accumulates (12).

A transcriptomic study of cold-shock in *Mycobacterium smegmatis* revealed distinct expression dynamics during the acclimation and adaptation phases. Alongside CspA and DEAD-box helicases, gene sets associated with cell wall remodelling, starvation, osmotic stress, and DNA topology were induced during acclimation, while ribosomal genes were down-regulated; both changes were reversed during adaptation to allow resumed growth (13).

In the present study, we analysed global transcriptional and translational changes in *S. coelicolor* A3(2) MT1110 following a sudden temperature shift from 30°C to 10°C under multiple growth conditions, providing a comprehensive view of cellular and metabolic processes perturbed during cold adaptation. Translational efficiencies (TE) were calculated to distinguish transcripts regulated at the transcriptional, translational, or both levels upon cold-shock. These data were further integrated into an established iModulon workflow to construct a systems-level landscape of the molecular networks governing the cold-shock response in *Streptomyces* (14).

## MATERIALS AND METHODS

### Establishment of cold-shock induction conditions and RNA extraction

Spores from the *S. coelicolor* prototrophic strain MT1110 were pregerminated in 100 mL of 2×YT medium for 7 hr at 30°C in an orbital shaker at 250 rpm (15, 16). The resulting germ tubes were used to inoculate 165 mL of SMM medium in 500 mL conical flasks equipped with stainless steel springs (approximately equal in length to the bottom circumference of the flask) at an initial inoculum concentration of approximately 5×10^5^ cfu/mL (16). Cultures were incubated at 30°C and 250 rpm until late exponential phase, corresponding to incubation time of either 17 hr 13 min (samples CT0A, CT0B, CS120A, CS120B, and CT120A) or 15 hr 45 min (samples CT0C, CT120B, CS120C, and CT120C). Immediately prior to cold-shock induction, a 10 mL aliquot of each culture was collected as a time=0 (t_0_) reference and stabilized by treatment with RNAProtect Bacteria Reagent (Qiagen, Hilden, Germany) at room temperature for 10 min. The resulting bacterial pellets were stored at -80°C for subsequent total RNA extraction. For polysome profiling, a separate 20 mL aliquot was harvested at the same timepoint to serve as the t=0 reference for monosome- and polysome-associated RNA fractions. The overall experimental design is schematically summarised in **Fig. 1**.

**Figure 1.**
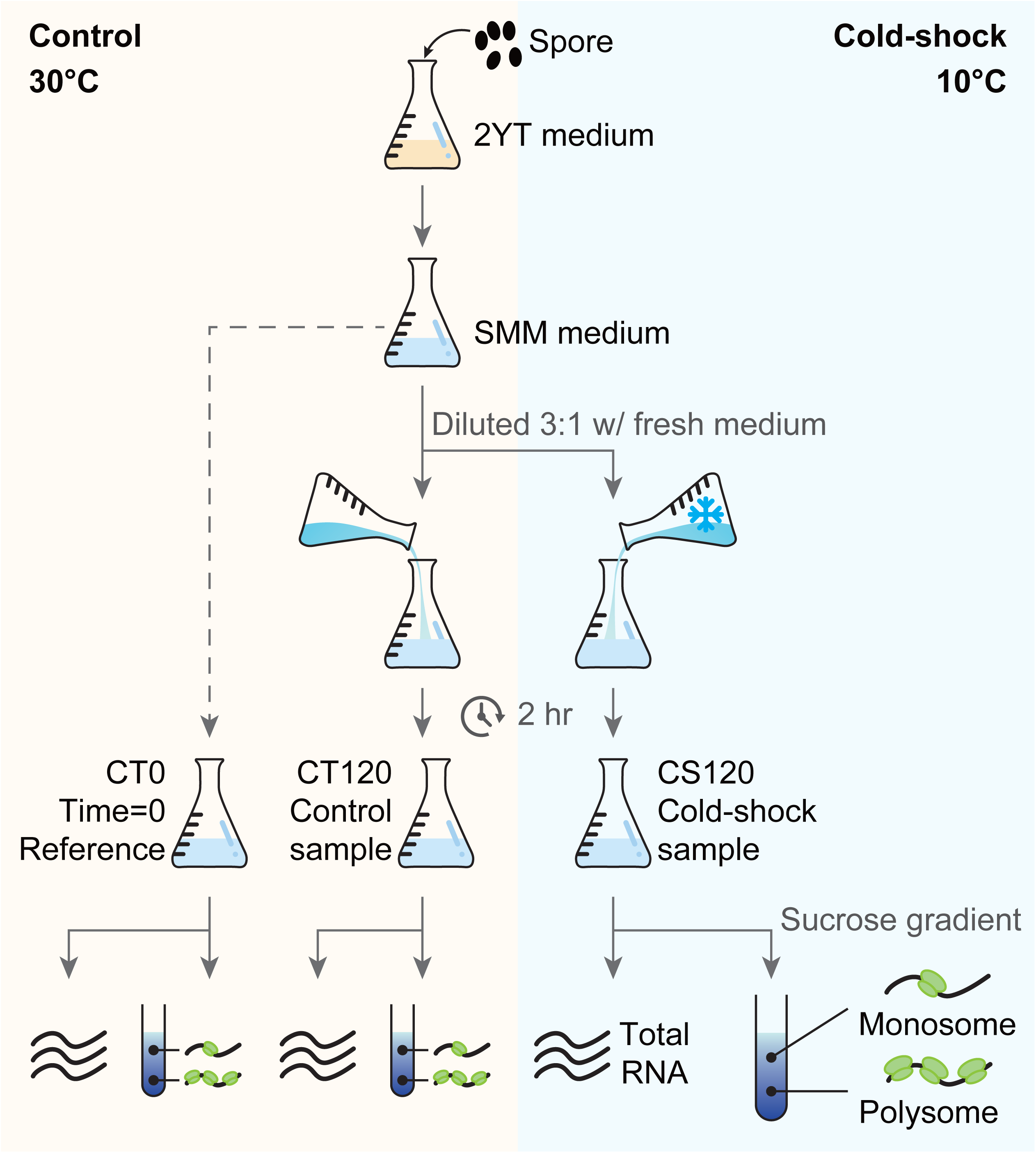
Schematic representation of the experimental design of the cold-shock experiments.

To induce cold-shock, 30 mL of late-exponential culture was rapidly cooled to 10°C by the addition of three volumes of fresh SMM medium pre-equilibrated at 4°C. In parallel, control cultures received three volumes of fresh SMM medium pre-equilibrated at 30°C to maintain the cultivation temperature. Following a 2 hr incubation under the respective conditions, a 40 mL aliquot of each culture was harvested for total RNA analysis and treated with RNAprotect as described above. Concurrently, a 50 mL aliquot was treated with chloramphenicol to arrest translation, and cells were lysed for downstream fractionation through sucrose density gradients. Two independent biological replicates were processed and analysed for all experimental conditions. Parallel cold-shock experiments were additionally conducted using surface-grown cultures on solid SMM agar, as well as liquid cultures cultivated in rich YEME medium. The resulting transcriptomic datasets were integrated into the global iModulon re-analysis reported herein.

### Polysome profiling

Polysome profiling was performed as previously described (1). To preserve the translational state of the cells at the time of harvest, chloramphenicol was added to designated culture aliquots to a final concentration of 80 µg/mL. Mycelia were immediately chilled on ice and harvested by centrifugation at 3,800×g for 4 min at 4°C. Mycelial pellets were resuspended in 3 mL of ice-cold lysis buffer (20 mM Tris-HCl [pH 8.0], 140 mM KCl, 1.5 mM MgCl_2_, 0.5 mM DTT, 1% Triton X-100, 80 μg/mL chloramphenicol, protease inhibitor cocktail (Sigma-Aldrich, St. Louis, MO), and 100 U/mL RNaseOUT (Thermo Fisher Scientific, Waltham, MA)). Mechanical cell disruption was achieved by vortexing in the presence of 5 mm glass beads four cycles of 20 sec each, with 1 min cooling intervals on ice between cycles. Mycelial debris was removed by centrifugation at 16,000×g for 5 min at 4°C, and the clarified supernatant was recovered for fractionation.

The supernatant was loaded onto a linear 10-50% sucrose density gradient prepared in gradient buffer (20 mM Tris-HCl [pH 8.0], 140 mM KCl, 5 mM MgCl_2_, 0.5 mM DTT, 80 μg/mL chloramphenicol) using a five-step dilution series from a 75% sucrose stock solution in nuclease-free water. Ultracentrifugation was performed at 166,880×g at 4°C. Following centrifugation, monosome and polysome fractions were resolved and collected using a Teledyne ISCO gradient fractionator coupled with a Foxy R1 fraction collector (Teledyne ISCO, Lincoln, NE). Collected fractions were immediately flash-frozen in liquid nitrogen and stored at -80°C.

To isolate polysome- and monosome-associated RNAs, three volumes of Trizol LS Reagent (Invitrogen, Waltham, MA) were added to 500 µL of each sucrose gradient fraction corresponding to the monosome or polysome peaks. The mixtures were homogenized by vortexing for 15 sec, incubated for 5 min at room temperature, and phase-separated by the addition of 200 µL chloroform, followed by an additional 15 sec vortex step and 5 min room temperature incubation. Following centrifugation, the aqueous upper phase was carefully recovered, and RNA was precipitated at -80°C for 1 h by the addition of 0.1 volumes of 3 M sodium acetate, 1 µL of glycoblue coprecipitant (Invitrogen), and 1 volume of isopropanol. RNA pellets were collected by centrifugation at 9,838×g for 1 h at 4°C, washed twice with 1 mL of ice-cold 75% ethanol, and resuspended in nuclease-free water.

### Transcriptome analysis

For total RNA extraction, bacterial pellets harvested from either 10 mL (t=0 reference) or 40 mL (cold-shock or control) cultures were resuspended in 200-400 µL of lysozyme solution (15 mg/mL in TE buffer) and incubated at room temperature for 15 min. Cells were subsequently lysed by the addition of 0.6-1.2 mL of RNeasy Lysis Buffer (Qiagen) supplemented with 10 µL/mL β-mercaptoethanol. The lysate was transferred to a 2 mL microfuge tube containing a 5 mm stainless steel bead (Qiagen) and homogenized using a Tissuelyser LT (Qiagen) at 30 Hz for 2 min. The homogenate was clarified by centrifugation at 16,627×g for 2 min, and the supernatant was recovered. Nucleic acids were purified by adding one volume of acid phenol:chloroform:isoamyl alcohol (25:24:1) (Fisher Scientific, Pittsburgh, PA), followed by vortexing for 30 sec and centrifugation at 16,627×g for 5 min at 4°C. This organic extraction was repeated twice more, followed by a final extraction with one volume of chloroform (16,627×g for 2 min at 4°C) to remove residual phenol. The final aqueous phase was processed using the miRNAeasy Mini Kit (Qiagen) following the manufacturer’s standard protocol. Total RNA was eluted in 30 µL of nuclease-free water and stored at -80°C. Residual genomic DNA was removed using the Turbo DNA-free Kit (Invitrogen).

Purified RNA from monosome/polysome fractions and total RNA preparations was quantified using a NanoDrop 1C spectrophotometer (Thermo Fisher Scientific) and quality-assessed using an Agilent 4200 TapeStation system (Agilent Technologies, Santa Clara, CA). RNA integrity number equivalent (RIN^e^) values for all analysed samples ranged from 6.5 and 9.2, with mean RIN^e^ values of 7.94 for RNA-seq libraries and 7.56 for polysome profiling fractions.

To enrich for mRNA, ribosomal RNA and highly abundant non-coding RNAs (*rnpB*, *ssrA*, and *srp*) were depleted from *S. coelicolor* total RNA using the NEBNext RNA Depletion Core Reagent Set (New England BioLabs, Ipswich, MA) in combination with custom antisense oligonucleotide probes (**Supplementary Data File S1**). These custom probes were developed following unsuccessful attempts to achieve efficient rRNA depletion using commercially available kits, specifically the Illumina Ribo-Zero Bacteria rRNA Depletion Kit and the Qiagen QIAseq FastSelect 5S/16S/23S Kit. Sequencing libraries were constructed using the NEBNext Ultra II Directional RNA Library Prep Kit for Illumina (New England BioLabs) and sequenced on an Illumina NextSeq 500 platform using a Mid Output v2.5 reagent kit (paired-end, 2×75 cycles (Illumina, San Diego, CA)).

### Data analysis

Sequencing data management and initial quality assessment were conducted via the Illumina BaseSpace web portal (http://basespace.illumina.com). Raw FASTQ files were analysed for quality metrics using FastQC (version 0.11.9) and aggregated using MultiQC (version 1.8) (17). The *S. coelicolor genome* assembly ASM20383v1 (RefSeq Accession No. GCF_00020385.1) was used as the reference genome. Transcript start and stop coordinates for all annotated open reading frames (ORFs) were extracted from the reference annotation and supplemented with empirical transcription start and stop sites previously mapped for *S. coelicolor* (18).

Read quantification was performed using Kallisto (version 0.46.2) against a transcriptome index derived from genome version GCF 000203835.1 ASM20383v1 (19). Transcript abundance estimates were aggregated to the gene level using tximport (version 1.20.0) (20). Downstream normalization and differential expression analysis were performed in R using the DESeq2 pipeline (21). Unsupervised sample clustering was evaluated using the R packages pheatmap and pcaMethods (22, 23). Differentially expressed genes (DEGs) within each RNA fraction (total RNA, polysome, or monosome) were identified using a stringent false discovery rate (FDR) threshold (*P_adj_* ≤ 0.05).

Significant changes in translational efficiency (TE) were determined using data from both the polysome and total RNA fractions, employing a generalized null hypothesis of:

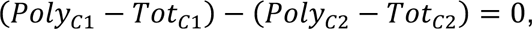

where *Poly* and *Tot* represent polysome and total RNA, respectively, and *C*1 and *C*2 represent the respective experimental conditions being compared. This model adheres to the fundamental definition of TE, whereby the TE of a given gene is defined as the ratio of its transcript abundance in the polysome fraction to its abundance in the total RNA pool. Based on the coordinated changes between the total transcriptome and polysome-associated fractions, genes were categorized into five distinct regulatory modes: **1) Forwarded** – Significantly altered in both the polysome and total RNA fractions, with no significant change in TE. **2) Exclusive** – Significantly altered in the polysome fraction only, with no significant change in the total RNA fraction. **3) Intensified (Potentiated)** – Significantly altered in the total RNA fraction, with a significant TE change in the same direction (both increased or both decreased). **4) Buffered** – Significantly altered in the total RNA fraction, with a significant TE change in the opposite direction. **5) None** – No significant change detected in either fraction (**Supplementary Data File S2**).

Functional annotation and gene set enrichment analyses were performed using clusterProfiler (24), with gene ontology (GO) annotations retrieved from the EMBL-EBI database (84.S_coelicolor.goa, July 2024). Protein-protein interaction and functional association networks were analysed using the STRING database (25). Conserved cis-regulatory DNA motifs within translation initiation regions (defined as -200 to +200 bp relative to the annotated start codon) of selected cold-shock regulon members were identified using the MEME Suite (26).

### iModulon analysis

The newly generated transcriptomic datasets were integrated into a previously compiled 469-sample *S. coelicolor* expression compendium and processed through an Independent Component Analysis (ICA) workflow (14). Briefly, expression profiles of the 7,710 core genes tracked in the baseline study were combined with the new cold-shock datasets. Transcripts Per Million (TPM) expression matrices from the cold-shock samples were Log_2_-transformed and baseline-subtracted using the mean expression values of the glucose-defined medium condition from the original compendium. iModulons were resolved from the expanded 505-sample compendium using the identical ICA pipeline constraints established in the original framework. The python core architecture used for computing and exploring iModulons is publicly available on GitHub via the modulome-workflow (https://github.com/avsastry/modulome-workflow) and pymodulon (https://github.com/SBRG/pymodulon) repositories.

## RESULTS AND DISCUSSION

### Transcriptional response to cold-shock in *Streptomyces coelicolor*

To determine the optimal conditions for cold-shock response, we collected samples cultured in SMM liquid medium at 15, 30, 60, and 120 min after shifting the temperature from 30°C to 10°C. A parallel set of control cultures was handled identically, but maintained at 30°C to control for the confounding effects of fresh medium addition (**Fig. 1**). Genes were defined as significantly differentially expressed (DEGs) in response to cold-shock only if they met a dual criterion: their expression was significantly altered two hours post-treatment (*Padj* < 0.05) compared to the matched 30°C control (10°C/120 min vs. 30°C/120 min), and they exhibited a significant change in the same direction (up- or down-regulation) relative to the pre-treatment baseline (10°C/120 min vs. 30°C/0 min) (**Fig. 2a**). This stringent filtering identified 811 cold-shock responsive DEGs, comprising 532 up-regulated (Cluster 1 in **Fig. 2b**) and 279 down-regulated genes (Cluster 2 in **Fig. 2b**). Notably, several hundred genes that were significantly altered in the 10°C/120 min vs. 30°C/120 min comparison were excluded from the cold-responsive dataset because they failed to show a significant change relative to the pre-treatment control (10°C/120 min vs. 30°C/0 min) (**Fig. 2a**). Given that cold-shock treatment inhibits growth, these excluded genes are likely growth-related transcripts whose expression changed more rapidly under optimal 30°C control conditions than during the decelerated growth at 10°C. It is worth noting that this experimental design evaluates expression based on steady-state transcript abundance, which reflects a combined effect of altered transcription rates (induction/repression) and changes in mRNA transcript half-lives.

**Figure 2.**
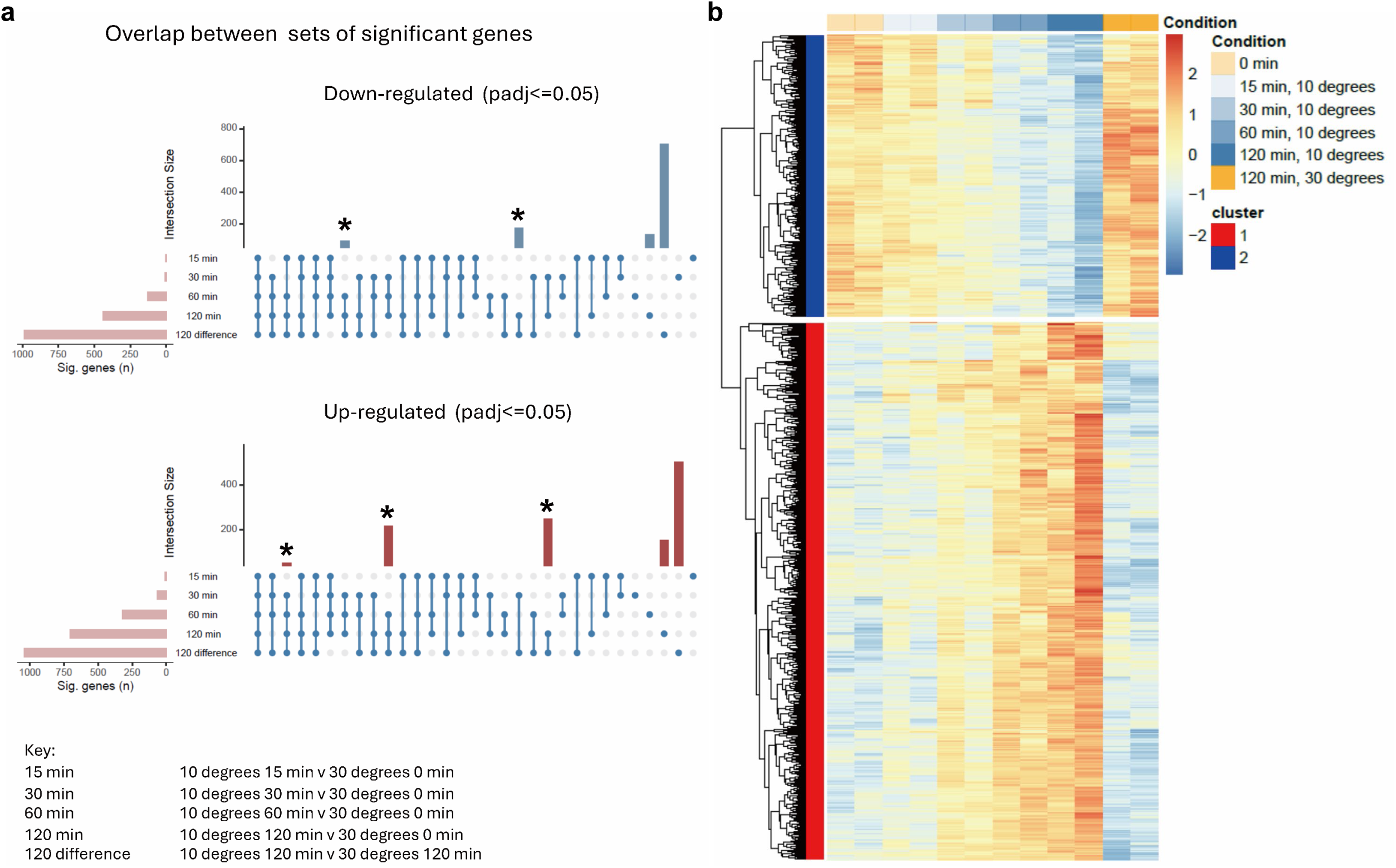
Identification of cold-shock induced genes. **(a)** Overlap between sets of significantly differentially expressed genes. Time course data from liquid SMM transcriptome experiment, Accession No. GSE225850, was used for analysis. Sets of genes that are significantly changed in the ‘120 difference’ group AND also significantly changed in any of the other groups that are relative to time 0 were taken forward in the analysis. These are considered ‘cold-shock’ responsive genes which are largely captured by the intersection of the differences identified in ‘10 degrees 120 min v 30 degrees 120 min’ and ‘10 degrees 120 min v 30 degrees 0 min’. These sets of genes are indicated by asterisks above the plots and represent the three conditions used in analysing translational control (Accession No. GSE229820). **(b)** Hierarchical clustering of cold-shock responsive genes. The 811 genes in the groups indicated by asterisks in Fig. 2a. The genes are detailed in **Supplementary Data File S3**.

Three CSP-encoding operons are massively induced following cold-shock. Although the *S. coelicolor* genome encodes a total of nine CSP homologues; only three were significantly differentially expressed following cold-shock, each located within one of three specific operons (**Fig. 3a**). Each of these operons encodes a highly conserved CSP (CspA homologue) alongside a ‘DEAD box’ RNA helicase, which is also massively induced by cold-shock. Additionally, a gene of unknown function and/or a gene encoding a cystathionine β-synthase (CBS) domain-containing protein are co-transcribed within the same operons. The expression profiles over time for the helicases adjacent to these cold-shock domain proteins are shown in **Fig. 3b**. All constituent genes of these three cold-shock induced CSP operons exhibited substantial up-regulation, reaching up to 1,973-fold (*SCO5920*) and 1,218-fold (*SCO4685*) induction (**Supplementary Data File S3**, **Supplementary Data File S4**).

**Figure 3.**
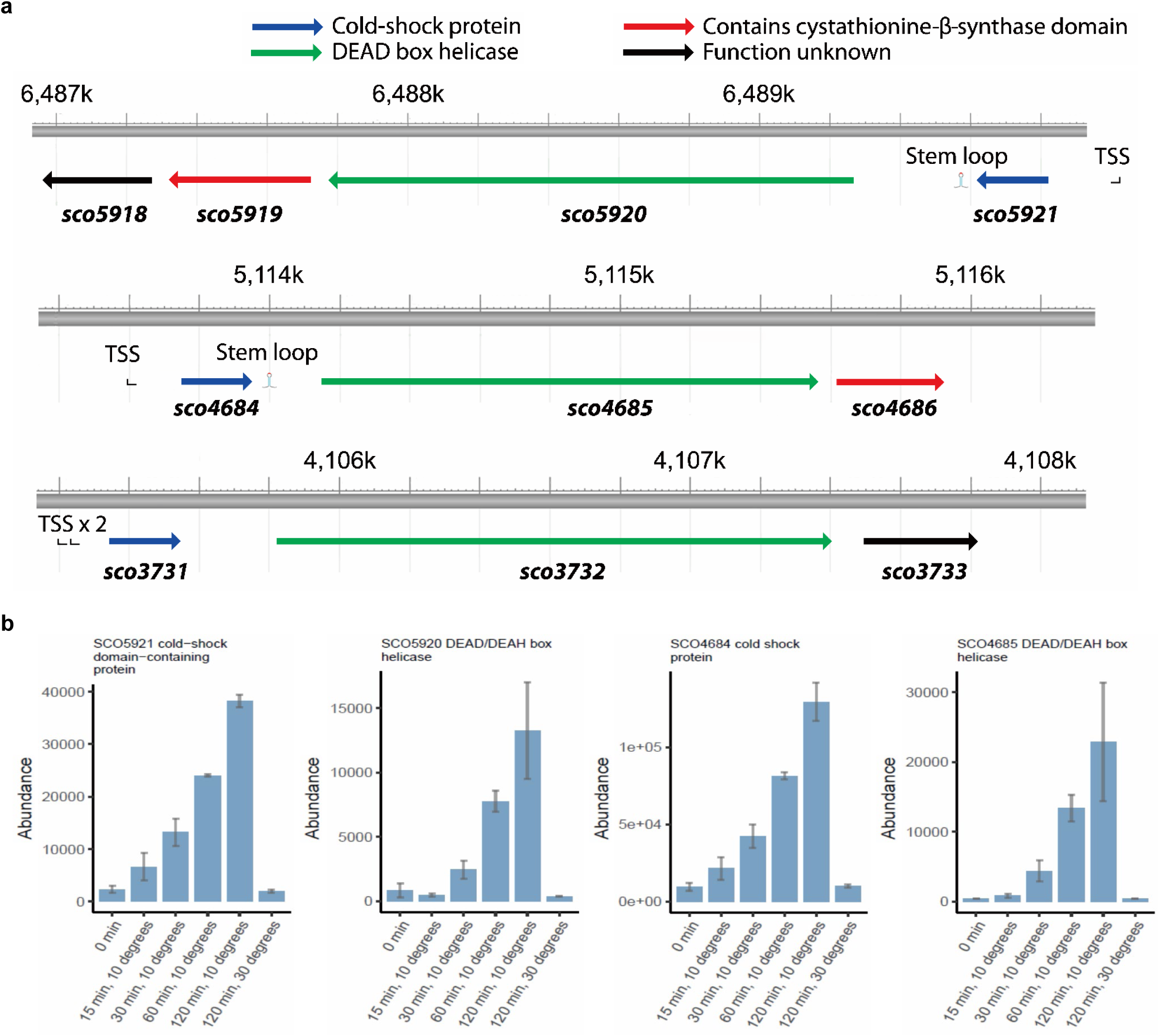
Analysis of major cold-shock protein operons. **(a)** Genetic organisation of the three major cold-shock induced operons of *Streptomyces coelicolor*. TSS indicates positions of known transcription start site(s), and in the two operons, *SCO5921-SCO5918* and *SCO4684-SCO4686*, the ‘cold-shock protein’ gene is immediately followed by a large, conserved, stem-loop structure (CS_IR_sco; Figure 7a). The scale above each diagram shows position on the chromosome (in kb). Adapted from images from NCBI Reference Sequence NC_003888.3 at www.ncbi.nlm.nih.gov. **(b**) Time course of cold-shock induced expression of genes encoding cold-shock domain proteins and helicases in the two major cold-shock operons.

### Functional enrichment analysis of the DEGs

Hierarchical clustering identified two principal clusters of DEGs corresponding to genes up- and down-regulated in response to cold-shock. Functional enrichment analysis of the 811 DEGs revealed no significant enrichment among the down-regulated genes (Cluster 2 in **Fig. 2b**). In contrast, four functionally enriched categories were identified among the cold-shock induced genes (Cluster 1 in **Fig. 2b**): 1) helicase activity (GO:0004386, GOMF); 2) transcription cis-regulatory region binding (GO:0000976, GOMF); 3) phenylalanine metabolism (sco00360, KEGG pathway); and 4) extracellular region (GO:0005576, GOCC) (**Fig. 4a** and **4b**).

**Figure 4.**
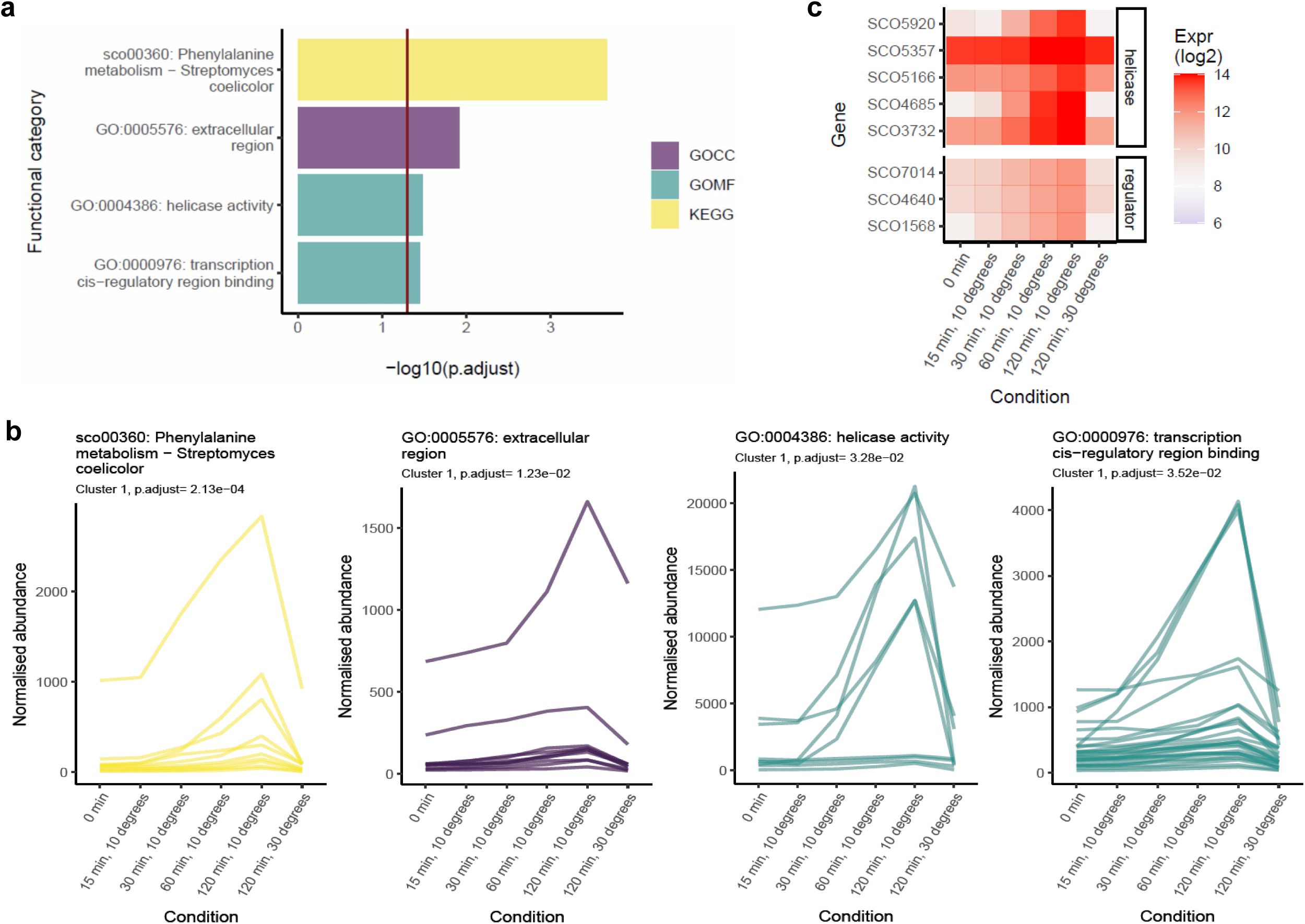
Analysis of cold-shock induced genes. **(a)** Functional enrichment analysis of the genes in Cluster 1 (Fig. 2b: heatmap). Key: GOCC; Gene Ontology Cellular Component; GOMF, Gene Ontology Molecular Function; KEGG, KEGG metabolic pathway. **(b)** Expression time course of the cold-shock induction of genes comprising the four significant functionally enriched groups. **(c)** Heatmap representing the time course of expression of the most highly expressed cold-shock induced genes encoding helicases and transcriptional regulators. The normalised expression values are given in **Supplementary Data File S5**.

The expression profiles of the nine helicase-encoding genes significantly up-regulated upon cold-shock are presented in **Fig. 4b**. This category includes three DEAD-box helicases encoded by *SCO4685*, *SCO5920* and *SCO3732*, each located within one of the three cold-shock operons (**Fig. 3a**). DEAD-box helicases are highly conserved cold-shock proteins across bacteria and function as global post-transcriptional regulators. By unwinding inhibitory mRNA secondary structures at low temperatures, they play a crucial role in restoring fundamental cellular processes disrupted by sudden cold exposure, including transcription and translation (27). This role is particularly important in *Streptomyces*, whose genome is characterized by an exceptionally high G+C content of approximately 73%, often exceeding 80% in coding sequences. The resulting propensity for stable secondary structure formation at low temperatures imposes a more severe obstacle to transcription and translation in *Streptomyces* than in other bacteria such as *E. coli* or *B. subtilis*. This functionally enriched category also includes two Rho transcription termination factors encoded by *SCO0537* and *SCO7051*. These RNA-DNA helicases facilitate transcription termination by binding nascent RNA and promoting its release from stalled transcription elongation complexes. This mechanism is particularly relevant in the context of cold-shock regulation, as rho-dependent transcription attenuation of cold-shock genes in *E. coli* has recently been identified as a key additional regulatory layer governing cold-shock gene expression (12). The expression profiles of the five most highly induced helicases are illustrated in **Fig. 4c**.

Among the transcriptional regulators, twenty-six were induced by cold-shock in *S. coelicolor* (**Fig. 4b**), of which three regulators encoded by *SCO1568*, *SCO4640*, and *SCO7014* displayed particularly pronounced induction profiles (**Fig. 4c**). Of these, only *SCO1568* has been functionally characterized; it encodes PqrA, a negative transcriptional autoregulator involved in paraquat resistance that co-regulates the expression of the downstream gene *pqrB* (28). *SCO7014* encodes a putative LacI-family transcriptional regulator, and *SCO4640* encodes a putative TetR-family transcriptional regulator; both remain functionally uncharacterized. In addition to these regulators, ten genes belong to the extracellular region category and were also induced upon cold-shock, the expression profiles of which are presented in **Fig. 4b** and listed in **Supplementary Data File S5**. The most highly induced gene in this group, *SCO3471*, encodes the DagA agarase precursor protein.

A further ten cold-shock induced genes are associated with the phenylacetic acid (PAA) degradation pathway, which represents an intermediate route in the catabolism of L-phenylalanine. PAA catabolism ultimately yields succinyl-CoA and acetyl CoA, both of which feed directly into the tricarboxylic acid (TCA) cycle. Notably, the PAA pathway has been reported to be highly regulated under diverse stress-associated conditions in *Acinetobacter* (29), suggesting a potential link between aromatic amino acid catabolism and stress signalling across bacterial genera. Collectively, these functional enrichment analyses reveal that the cold-shock transcriptional response in *S. coelicolor* is coordinated across multiple cellular levels, encompassing post-transcriptional regulation via RNA helicases, transcriptional reprogramming through regulatory proteins, remodelling of extracellular functions, and metabolic adaptation through amino acid catabolism. Together, these findings highlight the complexity and functional breadth of the cold-shock regulon in Streptomyces.

### Other highly induced cold-shock genes

Following cold-shock, the cell membrane undergoes a reduction in fluidity, which negatively impacts membrane polarization and the function of membrane-associated proteins (30). The gene *des* (*SCO3682*), homologous to the *B. subtilis* delta fatty acid desaturase-encoding gene, acts to restore membrane fluidity by introducing double bonds into membrane-bound fatty acids. *SCO3682* is strongly induced upon cold-shock, along with *SCO3681* (the first gene of the operon), *SCO3684*, and *SCO3685*, the latter three of which encode proteins of currently unknown function. Cold-shock also induces the expression of two enzymes involved in cell wall and cell wall glycan biosynthesis: *SCO7050* and *SCO6179*. *SCO7050* encodes a D-alanyl-D-alanine carboxypeptidase, a member of the penicillin-binding proteins (PBPs) family, which is inhibited by β-lactam antibiotics and is involved in peptidoglycan synthesis and remodelling. *SCO6179* encodes a nucleotide sugar dehydratase and constitutes the first gene of the *cwg* operon, which encodes enzymes involved in cell wall glycan biosynthesis (31).

Cold-shock also induced the expression of several genes encoding efflux pumps and membrane transporters. These includes *SCO4120,* which encodes an integral membrane protein with homology to multidrug resistance proteins that confer resistance to ciprofloxacin and chloramphenicol and enhance oxidative stress tolerance (32), *SCO3366*, which encodes a rifampicin efflux pump (33), *prqB* (*SCO1567*), which encodes a Major Facilitator Superfamily (MFS) efflux pump conferring resistance to the redox-cycling agent paraquat (28), and *cmlR2* (*SCO7662*), which mediates chloramphenicol resistance (34). Additional cold-shock induced genes include *SCO4641*, an MFS superfamily transmembrane efflux pump, as well as the first two genes of the *areABCD* gene cluster (*SCO3956 – SCO3959*) which encode two distinct ABC transporter systems. Within this cluster, *SCO3956* is universally conserved across all domains of life, whereas *SCO3959* is conserved exclusively within the bacterial domain.

### Identification of conserved motifs in the 5′-regions of cold shock genes

The observation that three transcriptional regulators (TFs) are strongly induced by cold-shock in the SMM liquid datasets (**Fig. 4c**) prompted a search for genes displaying correlated expression profiles, with the aim of identifying potential targets of these TFs (**Supplementary Data File S6**). High Spearman correlation coefficients were observed with several genes, including the CspA-encoding homologues, *SCO4684* and *SCO5921*. Two small RNAs, sRNA126 and sRNA208, also ranked among the top correlated genes. Subsequent searches for conserved DNA sequence motifs were conducted using the MEME suite (**Supplementary Fig. S1a-S1d**) (26). The most prominent conserved motif identified was a stem-loop structure located downstream of the two major cold-shock genes, *SCO5920* and *SCO4685* (**Supplementary Fig. S1a** and **Fig. 5**).

**Figure 5.**
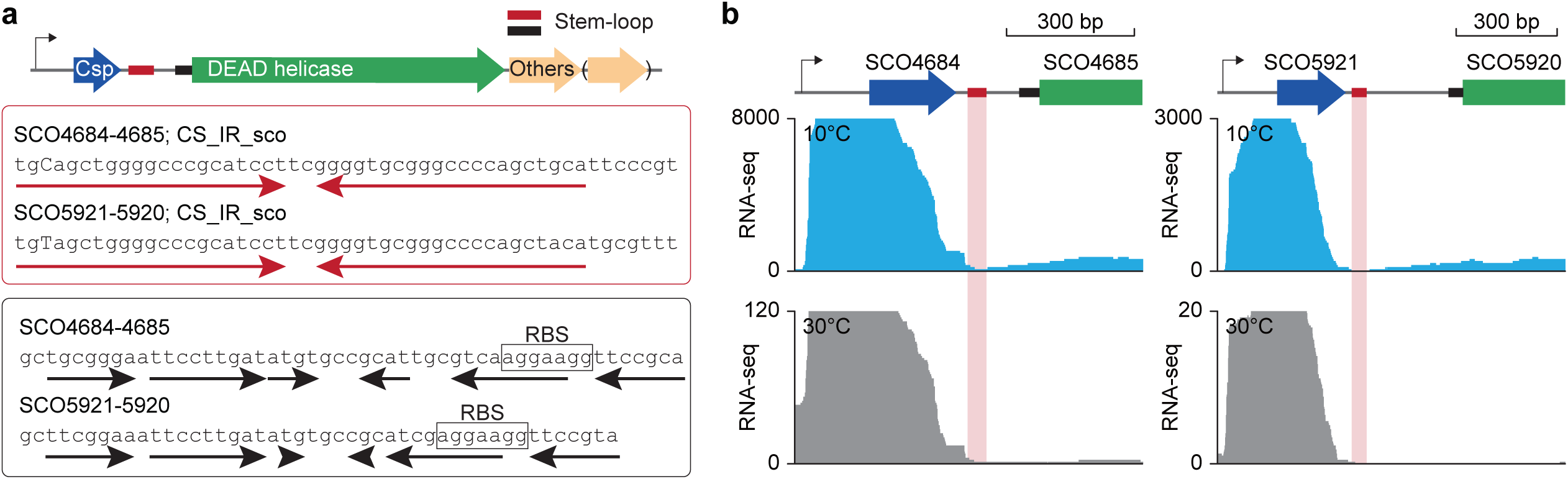
Analysis of the intergenic regions between the *csp* and DEAD-box RNA helicase-encoding genes in the major cold-shock operons. (**a**) Probable RNA stem-loop structures identified in the intergenic region between *csp* and DEAD helicase. (**b**) RNA-seq read profile of *csp* (SCO4684 and SCO5921) and downstream DEAD helicase encoding (SCO4685 and SCO5920) loci, from cold-shocked and control (30°C) cultures.

### iModulon-based analysis of cold-shock response in *Streptomyces coelicolor* A3(2)

To further characterize the global transcriptional response to cold-shock, an iModulon-based analysis was performed to identify a higher-order regulatory architecture complementing the insights derived from conventional DEG analysis (35). An iModulon is defined as a group of co-transcriptionally regulated genes resolved through Independent Component Analysis (ICA) of transcriptomic datasets, frequently overlapping with the regulon of a specific transcriptional regulator and representing a discrete cellular function. When the cold-shock transcriptomic data were initially interpreted using previously established iModulons, only approximately 60% of transcriptomic variance could be explained (**Supplementary Fig. S2**) (14). Given that the prior iModulon framework was constructed from datasets lacking cold-shock conditions, we hypothesized that integrating the new cold-shock transcriptomes into the ICA workflow would reveal novel gene modules serving as key components of the cold-shock response in *S. coelicolor*.

Accordingly, ICA was applied to an expanded compendium comprising 505 transcriptomic profiles, encompassing the previously compiled 469 datasets and 36 newly generated cold-shock transcriptomes. This analysis revealed a total of 129 iModulons (**Supplementary Data File S7-10**), representing an increase of 12 iModulons compared to the previous study (**Supplementary Fig. S3**) (14). Of these, 19 iModulons were newly identified and associate with diverse functions (**Fig. 6a** and **Supplementary Fig. S3**). Notably, the original cold-shock iModulon, which previously encompasses genes within the *cspA* operons, was resolved into two distinct iModulons with differing genetic contexts: the ‘cold-shock regulatory’ iModulon, comprising upstream elements of the *csp* operons (including *csp* and DEAD-box helicase-encoding genes) together with five transcriptional regulators, and the ‘*csp* downstream’ iModulon, which lacks *csp* but contains the DEAD-box helicase genes and other downstream genes of unknown functions (**Fig. 6b** and **Supplementary Table S2**).

**Figure 6.**
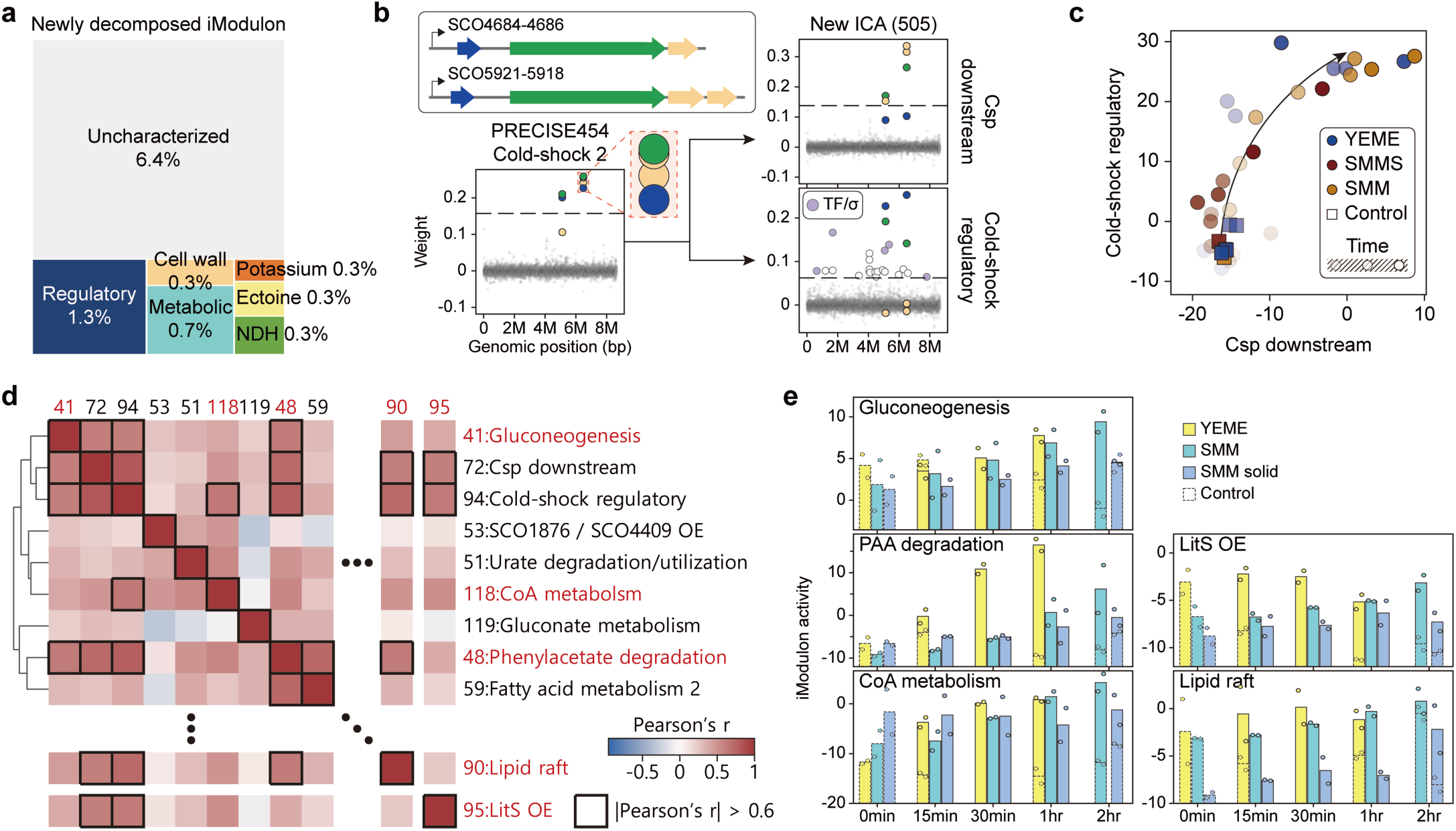
Overview of iModulon-based analysis on cold-shock response in *S. coelicolor*. **(a)** Functional categorization of the 19 newly emerged iModulons. **(b)** The original cold-shock iModulon split into two iModulons with distinct genetic contexts. The cold-shock regulatory iModulon includes upstream elements of *csp* operons (*csp* and DEAD helicase) along with 5 transcriptional regulators, while the *csp* downstream iModulon lacks *csp* and contains the DEAD helicase and the other downstream genes of unknown functions. **(c)** Activity relationship between the two *csp*-related iModulons. The regulatory iModulon is activated earlier, followed by the downstream iModulon. Arrow indicates progression over time after cold-shock. **(d)** iModulons associated with the cold-shock response. Hierarchical clustering of iModulons was performed based on the Pearson correlation coefficients of iModulon activities. iModulons clustered in the vicinity of the two cold-shock iModulons are visualized, together with other iModulons with high activity correlation (Pearson’s r > 0.6). **(e)** Activity of iModulons with high activity correlation (Pearson’s r > 0.6) against cold-shock iModulons.

Among the 26 cold-shock-induced transcriptional regulators identified as DEGs, five were associated with the cold-shock regulatory iModulon, including four transcription factors (*SCO1145*, *SCO1568*, *SCO4640*, and *SCO7014)* and one ECF sigma factor (*SCO4908*; SigQ), strongly suggesting their direct involvement in the cold-shock response (**Supplementary Table S2**). Although the precise regulatory roles of these transcription factors remain to be elucidated, SigQ has been previously characterized as a pleiotropic regulator involved in environmental adaptation and secondary metabolism (36), and its association with the cold-shock regulatory iModulon further underscores its specific role in the cold-shock response. A palindromic sequence motif was additionally identified within the promoter regions of the cold-shock regulatory iModulon genes, implicating it as a potential cis-regulatory element mediating this response (**Supplementary Fig. S4**).

Remarkably, the two cold-shock iModulons are activated sequentially, with activation of the ‘cold-shock regulatory’ iModulon preceding that of the ‘*csp* downstream’ iModulon (**Fig. 6c**). To investigate the regulatory mechanism underlying this sequential activation, the intergenic regions between the *csp* genes and DEAD-box helicase-encoding genes were examined. This analysis led to the prediction of two independent RNA stem-loop secondary structures located downstream of the primary cold-shock genes *SCO4684* and *SCO5921* (**Fig. 5a**). One stem-loop, immediately downstream of these *csp* genes, was also identified by MEME Suite analysis (**Supplementary Fig. S1a**) and shares sequence similarity with a *Streptomyces-*specific transcription termination motif previously described in other species (37–40). Consistent with a functional transcription termination role, RNA-seq read coverage showed a marked decrease immediately downstream of this motif (**Fig. 5b**).

The second stem-loop structure spans the RBS sequence of the downstream DEAD-box helicase-encoding genes, suggesting a potential role in translational regulation mediated by these downstream elements. Taken together with the observed cold-shock induction of the Rho-dependent transcription termination factors, *SCO0537* and *SCO7051*, these structures may represent transcriptional regulatory switches constituting an additional layer of cold-shock gene regulation, analogous to mechanisms described in *E. coli* (12). Moreover, the five transcriptional regulators identified within the cold-shock regulatory iModulon may drive the transcription of downstream genes through a regulatory cascade, while transcription of the upstream *csp* genes terminates at the preceding terminator motif. Overall, this iModulon-based analysis provides critical insights into the complex multi-layered regulatory architecture governing the cold-shock response in *S. coelicolor*.

### iModulon activities analysis

To identify metabolic pathways closely associated with the cold-shock response, pairwise hierarchical clustering of iModulon activity correlations was performed across all cold-shock samples (**Supplementary Fig. S5**). Seven iModulons clustered in close proximity to the two cold-shock iModulons, of which three iModulons, gluconeogenesis, CoA metabolism, and phenylacetate degradation (PAA), showed high activity correlation with the cold-shock modules (Pearson’s r > 0.6), suggesting the coordinated involvement of these metabolic pathways in the cold-shock response (**Fig. 6d** and **e**).

Specifically, gluconeogenesis provides the carbon precursors required for trehalose biosynthesis, a disaccharide that functions as an essential cryoprotectant by stabilizing membranes and proteins during abiotic stress, including cold exposure (41, 42). Concurrently, coenzyme A (CoA) metabolism is implicated in cold-shock adaptation in *Streptomyces* through the reconfiguration of membrane fatty acid composition and the homeostatic regulation of metabolic flux necessary for survival at low temperatures. The physiological importance of CoA is underscored by its indispensable role in fatty acid synthesis and the modulation of energy metabolism under stress conditions. Furthermore, the PAA pathway catabolizes aromatic compounds to yield succinyl-CoA and acetyl-CoA, both of which serve as direct substrates for the tricarboxylic acid (TCA) cycle (43). Collectively, the coordinated induction of these pathways alongside the two cold-shock iModulons indicates that they jointly supply the energy and biosynthetic intermediates required to sustain essential cellular function at low temperatures.

Two additional iModulons related to lipid raft reorganization and *litS* overexpression also showed high activity correlation with the cold-shock iModulons (Pearson’s r > 0.6) (**Fig. 6d**). Cold exposure is well established to induce the lateral redistribution and coalescence of lipid rafts into larger functional microdomains, which serve as specialized, more ordered regions within the bacterial membrane. Bacteria exploit these functional membrane microdomains (FMMs) to concentrate specific sensor kinases and stress-response proteins, effectively compartmentalizing them from the bulk fluid membrane to facilitate efficient stress signalling (44). Meanwhile, the ECF sigma factor LitS is known to regulate carotenoid biosynthesis (45), and the accumulation of carotenoids has been associated with enhanced resistance to freeze-thaw stress (46). However, as the LitS regulon is predominantly activated by light exposure, the possibility that the observed *litS* induction resulted from incidental daylight exposure during the cold-shock experiment cannot be excluded at present.

### Post-transcriptional regulation of the cold-shock response

The rate-limiting step in protein synthesis is translation initiation, and thus ribosome attachment to transcript serves as a reliable proxy for translational activity (47). To explore translation-level dynamics in response to cold-shock, cultures were subjected to concurrent RNA-seq and polysome profiling following 120 min of cold-shock treatment at 10°C (GSE229820). Translational Efficiency (TE) is defined as the ratio of polysome-associated transcript abundance to total transcript abundance for a given gene, such that significant alterations in TE are indicative of post-transcriptional regulation. TE changes were assessed using the *deltaTE* protocol (48), classifying genes into four distinct regulatory groups, including *forward*, *exclusive*, *intensified* (or potentiated), and *buffered*. Overall, a strong positive correlation was observed between changes in total RNA and polysome-associated fractions across all comparisons (10°C/120 min vs. 30°C/0 min, R = 0.839; 10°C/120 min vs. 30°C/120 min, R = 0.835; 30°C/120 min vs. 30°C/0 min, R = 0.799; all *P* < 2.2×10⁻¹⁶) (**Fig. 7**; **Supplementary Data File S2**).

**Figure 7.**
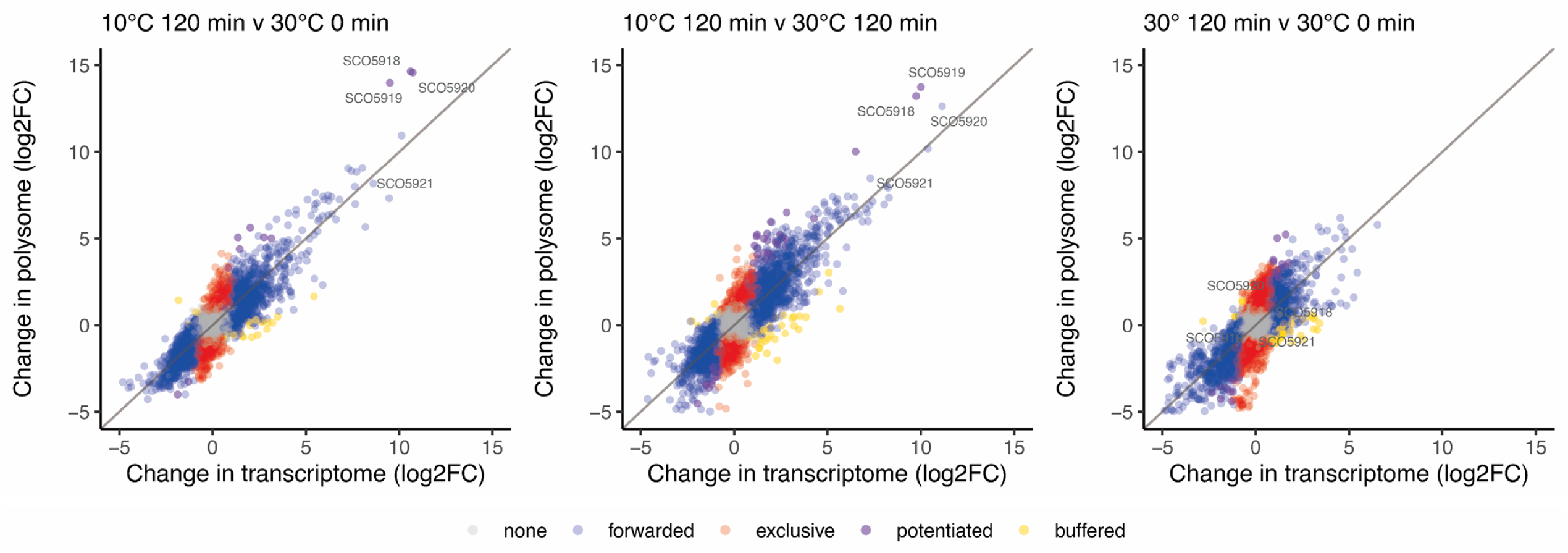
Scatter plots of the transcriptome data relative to the polysome profiling data. There are good correlations between changes in relative abundance of transcripts in the polysome fraction compared to their respective changes in abundance in the transcriptome: ‘10°C 120 min v 30°C 0 min’, R=0.839 (p-value < 2.2e-16); ‘10°C 120 min v 30°C 120 min’, R=0.835 (p-value < 2.2e-16); ‘30° 120 min v 30°C 0 min’, R=0.799 (p-value < 2.2e-16)). The different classifications of expression, forwarded, exclusive, potentiated and buffered are colour-coded below the figure. The key potentiated cold-shock operon gene products, SCO5921-SCO5918, are indicated on the plots. The data used to generate these plots is given in Supplementary Data File S2.

The *forward* group comprises genes regulated primarily at the transcriptional level, whose TE remains unchanged relative to the reference condition, indicating translation in direct proportion to transcription.

The *exclusive* group encompasses genes regulated solely at the translational level, with no significant change in total transcript abundance but significant up- or down-regulation in ribosome-associated fractions. Functional enrichment analysis of the exclusively enhanced group identified significant enrichment of protein phosphorylation (GO:0006468; FDR = 0.0338), consistent with the requirement for rapid post-transcriptional signalling following sudden temperature downshift. Conversely, the exclusively repressed group includes the glutamine synthetase-encoding gene *SCO2198* and components of the glycine cleavage system, with enrichment in oxidative phosphorylation (*SCO00190*, FDR = 1.68 e-05), metabolic pathways (*SCO01100*, FDR = 3.17 e-10), biosynthesis of secondary metabolites (*SCO01110*, FDR = 6.19 e-06), carbon metabolism (*SCO01200*, FDR = 0.00029), and amino acid biosynthesis (*SCO01230*, FDR = 0.00067) **(Supplementary Data File S2, Supplementary Fig. S6)** (49).

Among the exclusively enhanced transcripts, *SCO4124* and *SCO2216* encode homologues of the *B. subtilis* DesK sensor kinase and DesR response regulator, respectively, which together constitute a membrane fluidity-sensing system activated via DesR phosphorylation (50). *SCO3956* and *SCO3959*, the terminal genes of a four-gene operon encoding an ABC transporter complex involved in S-adenosylmethionine (SAM) signal transduction (51), are also exclusively enhanced: notably, the first two genes of this operon are additionally induced at the transcriptional level, indicating multi-layered activation.

The *intensified* (potentiated) group contains genes regulated at both transcriptional and translational levels in the same direction. Functional enrichment analysis revealed over-representation of ABC transporters and integral membrane proteins among potentiated-up genes, while potentiated-down genes were enriched in respiratory chain components, cytochrome c oxidase activity, and nitrate metabolic processes (**Supplementary Data File S2**). With the exception of *SCO5921*, the remaining three genes of the major cold-shock operon were classified as potentiated, underscoring their crucial role in the early cold-shock response. Other notable potentiated-up transcripts include *SCO4640*, a highly induced transcription factor implicated in downstream adaptive responses (**Fig. 4c**).

The three operonic genes, *SCO4584-SCO4586* are all potentiated and encode a functional ABC transporter complex. Additionally, the ‘potentiated-up’ cluster *SCO7748*-*SCO7750* encodes hypothetical proteins, where *SCO7750* specifies a putative anti-anti sigma factor and *SCO7748* encodes an RsbU-homologous serine phosphatase; *SCO7748* and *SCO7750* represent a highly conserved gene pair across prokaryotes. Importantly, *B. subtilis* RsbU forms an integral part of the ‘stressosome’ complex, where RsbR acts as a dual positive and negative regulator of sigma-B activity (52). Other highly induced transcripts include *SCO7050* (‘potentiated up’), which encodes a D-Ala-D-Ala carboxypeptidase and *SCO3830* (BkdB2) (‘potentiated up’), which encodes a probable branched chain alpha keto acid dehydrogenase subunit, and *SCO2927*, which encodes a putative 4-hydroxyphenylpyruvate dioxygenase (HppD). HppD is central to tyrosine catabolism, yielding fumarate and acetoacetate to feed the TCA cycle and provide essential nitrogen, carbon, and energy substrates for survival (53).

The *buffered* group contains genes regulated at both levels but in opposing directions. Several buffered genes are directly involved in translation, including *SCO7600*, encoding a putative alanyl-tRNA synthetase (AlaS2), which is transcriptionally enhanced but translationally repressed and is a known component of the WblC regulon governing translation-related gene expression under antibiotic stress (54). *SCO6149*, encoding an ATP/GTP-binding protein involved in 30S ribosomal subunit maturation. *SCO1321*, encoding translation elongation factor Tu-3, and *cmlR2* (*SCO7662*), a chloramphenicol resistance gene that is transcriptionally induced but translationally repressed.

## CONCLUSIONS

In this study we have established the experimental conditions required to elicit a robust cold-shock response in *Streptomyces*. In contrast to the streptomycete heat-shock response, the cold-shock response is principally regulated at the transcriptional level. Functional enrichment analysis of the initial transcriptome experiments conducted in liquid SMM cultures identified four principal enriched categories, including transcription regulation, helicase activity, phenylalanine metabolism, and extracellular functions. The critical importance of RNA helicase activity in destabilizing inhibitory mRNA secondary structures, thereby sustaining fundamental cellular processes at low temperatures, is particularly pronounced in streptomycetes, whose protein-coding sequences are characterized by an exceptionally high G+C content, exceeding 80%.

By combining all cold-shock transcriptomic datasets across diverse culture conditions, an iModulon-based analysis was performed to provide a system-level view of the *Streptomyces* transcriptional regulatory network responding to thermal stress. This analysis revealed two sequentially activated cold-shock iModulons whose successive expression is governed by cis-acting regulatory elements located downstream of the principal CspA-encoding homologues, *SCO4684* and *SCO5921*. Five additional iModulons were identified whose activities closely correlated with those of the two-core cold-shock iModulons, providing mechanistic insights into the coordinated metabolic and structural rearrangements that mitigate cold-induced cellular damage, encompassing gluconeogenesis, CoA metabolism, phenylacetate degradation, and lipid raft remodelling.

At the molecular level, three cold-shock operons encoding CspA homologues and DEAD-box helicases were massively induced. Five cold-inducible transcription factors were identified, four of which (SCO1145, SCO1568, SCO4640 and SCO7014) were previously uncharacterized, while SCO4908 (SigQ) is a known regulator, likely to direct the metabolic transition between primary and secondary metabolism (36).

Highly conserved sequence motifs were identified upstream of the three cold-shock induced operons, while prominent stem-loop structures were predicted downstream of the *cspA*-encoding homologues, *SCO4684* and *SCO5921*. Future biochemical studies are warranted to precisely dissect the respective contributions of these elements to the transcriptional and translational control of the cold-shock response. Notably, the most strikingly potentiated genes reside within the *SCO5921*-*SCO5918* operon, underscoring its indispensable role in physiological acclimation. Given that the helicase-encoding genes within this locus are transcriptionally induced up to 2,000-fold, the tightly efficient regulatory mechanisms governing this extraordinary scale of induction hold considerable promise for future synthetic biology applications.

Finally, this study has uncovered novel genetic components integral to the *Streptomyces* cold-shock response. For example, the newly identified gene pair, *SCO0818* and *SCO0819*, which encodes an ABC transporter ATP-binding protein and a cognate transmembrane protein, respectively, is predicted to fulfil a crucial, as-yet-unexplored physiological role in cold-shock adaptation.

## Supporting information

Transcriptome and polysome data

Main transcriptome clusters

Gene expression matrix

Main regulated pathways from SMM data

Genes correlating with the three CS TFs

iModulon X spreadsheet

iModulon M spreadsheet - placeholder

iModulon A spreadsheet

iModulon list spreadsheet

## DATA AVAILABILITY

*The gene expression data underpinning this article are available from* NCBI’s Gene Expression Omnibus at https://www.ncbi.nlm.nih.gov/geo *and can be accessed with the following accession numbers:* GSE225850, GSE229820, GSE220990.

## AUTHOR CONTRIBUTIONS

R.T.E. Data curation, Formal analysis, Investigation, Methodology, Software, Visualisation, Funding acquisition. Y.L. Data curation, Formal analysis, Investigation, Methodology, Software, Visualisation, Writing – original draft, Writing – review & editing. A.H. Data curation, Formal analysis, Methodology, Software, Supervision, Validation, Visualisation, Writing – original draft. B.-K.C. Formal analysis, Methodology, Project administration, Resources, Software, Supervision, Validation, Visualization, Writing – original draft, Writing – review & editing, Funding acquisition. G.B. Conceptualisation, Methodology, Project administration, Supervision, Validation, Writing – original draft, Writing – review & editing. C.P.S. Conceptualisation, Formal analysis, Methodology, Project administration, Resources, Supervision, Validation, Visualization, Writing – original draft, Writing – review & editing, Funding acquisition.

## ACKNOWLEDGEMENTS

We are grateful for technical advice and support from Dr Kenji Nakahigashi and Prof Hirotada Mori at NAIST (Japan) and Dr Rocio Martinez-Nunez from KCL (UK) and thank Prof Gilles van Wezel from Leiden University for helpful discussions.

## FUNDING

This work was generously supported by the philanthropists, Michael and Maureen Chowen, and by the University of Brighton, who provided a PhD scholarship (to RTE) and was also supported by the National Research Foundation (NRF) of Korea funded by the Korean Government (grant no. RS-2024-00352229 and RS-2025-02216377 to B-KC) and the KAIST Strategic Research Program (N10260074 to B-KC). RTE also contributed financially to research costs.

## CONFLICT OF INTEREST STATEMENT

None declared.

## SUPPLEMENTARY FILES

### SUPPLEMENTARY TABLES

**Supplementary Table S1**. **Conserved sequence motifs in cold shock induced genes.** The consensus sequences of each motif identified in this study are presented together with the coordinates of each hit and their respective statistical significance (generated using MEME).

**Supplementary Table S2. Gene membership of the ‘cold-shock regulatory’ iModulon.**

### SUPPLEMENTARY DATA FILES

**Supplementary Data File S1**: Custom oligonucleotides designed in this study for depleting rRNAs and selected non-coding RNAs. Filename: Supplementary Data File S1 (Word format).

**Note**: for each Excel spreadsheet a README sheet is included with an explanatory key for each column.

**Supplementary Data File S2**: p.GSE229820.combined.rna_v_ps.xlsx

**Supplementary Data File S3**: p.Heatmap.GSE225850_clusters.xlsx

**Supplementary Data File S4**: DESeq2_SMM_GSE225850_GSE229820.xlsx

**Supplementary Data File S5:** p.ORA.genes.GSE225850.xlsx

**Supplementary Data File S6**: Genes correlating with the three CS TFs GSE225850_reg_correlations.xlsx

**Supplementary Data File S7**: iModulon_X.xlsx

**Supplementary Data File S8**: iModulon_M.xlsx

**Supplementary Data File S9**: iModulon_A.xlsx

**Supplementary Data File S10**: iModulon_list.xlsx

### SUPPLEMENTARY FIGURES

**Supplementary Figure S1a-S1d**: Sequence motif identification. See Supplementary Figure S1a-S1d file for full text description of the figures.

**Supplementary Figure S2:** The percentage of transcriptomic variance explained by previously computed iModulons.

**Supplementary Figure S3:** Comparison between previously computed iModulons and newly computed iModulons with cold-shock conditions. To identify newly emerged or lost modules from the integration of cold shock conditions, cosine similarity between old and new iModulons was calculated; iModulon pairs with cosine similarity > 0.2 were linked together, indicating that the old iModulon has been retained, and the unlinked iModulons were regarded as lost or newly emerged modules. If multiple iModulons showed cosine similarity > 0.2 to a single original iModulon, they were considered as split modules.

**Supplementary Figure S4:** Sequence motif identified in the promoter regions of cold-shock regulatory iModulon genes. Motif search was performed using the 50-nt upstream sequences from the transcription start sites reported by Jeong et al. with the MEME Suite using the ZOOPS option.

**Supplementary Figure S5:** Hierarchical clustering of iModulon activities across cold-shock samples. The black square indicates the subcluster containing the two cold-shock iModulons.

**Supplementary Figure S6:** Pathways downregulated following cold-shock. (a) KEGG pathways; (b) Biological Process (Gene Ontology) enrichment.

## Notes

### Competing Interest Statement

The authors have declared no competing interest.

### Summary of Updates

After the original manuscript was submitted the authors became aware of an innovative unsupervised machine learning approach for analysing the structure of transcriptomes, called iModulon, to identify all co-ordinately-controlled 'modulons' within a biological system. Our collaborators, led by Byung-Kwan Cho recently published an iModulon-based model in the antibiotic-producing bacterium, Streptomyces coelicolor. However, they had not incorporated transcriptome data that pertained to the cold-shock response because it hadn't existed at the time. We invited the lead author of the iModulon paper and Prof Cho to investigate all our raw transcriptome and translatome cold-shock data, to integrate into their Streptomyces 'iModulon' system. The insights from this new analysis have significantly extended our understanding of how Streptomyces remodels its metabolism and cellular organisation in response to a rapid drop in temperature and strongly justified inclusion of these results along with our previous results and their inclusion as co-authors. The study now adds substantially to our knowledge of how complex high %G+C-content bacteria tackle the physico-chemical shock of a temperature downshift in order to survive such extreme stress.

