## Supplementary Figures and Tables for "Global analysis of the cold-shock response in the model antibiotic producing actinomycete, *Streptomyces coelicolor* A3(2)"

Supplementary Material (Figures and Tables) and Supplementary Data File S1 for
### Global analysis of the cold-shock response in the model antibiotic producing actinomycete, *Streptomyces coelicolor* A3(2)

R. Tony Evans[1#], Yongjae Lee[2,3#], Andrew Hesketh[1], Byung-Kwan Cho[2,3,*], Colin P. Smith[1,4,5,*] and Giselda Bucca[1,4,*]

[1]School of Applied Sciences, University of Brighton, Lewes Road, Brighton, UK
[2]Graduate School of Engineering Biology, Korea Advanced Institute of Science and Technology, Daejeon 34141, Republic of Korea
[3]KI for the Bioinnovation, Korea Advanced Institute of Science and Technology, Daejeon 34141, Republic of Korea
[4]Present address: School of Immunology & Microbial Sciences, Faculty of Life Sciences & Medicine, Guy's Hospital, Borough Wing, Great Maze Pond, King's College London, London, SE1 9RT, UK
[5]Present address: School of Biosciences, Faculty of Health and Medical Sciences, University of Surrey, Surrey, GU2 7XH, UK.

[#]These authors contributed equally.

*Corresponding authors: Byung-Kwan Cho (, ORCID: https://orcid.org/0000-0003-4788-4184), Colin P. Smith (, ORCID: https://orcid.org/0000-0001-9042-2263), Giselda Bucca (, ORCID: https://orcid.org/0000-0001-5286-7771)

[†]This paper is dedicated to the memory of Professor David A. Hodgson.

**Contents**:
Supplementary Figures: 1 to 6
Supplementary Tables: 1 to 2
Supplementary Data File S1

Supplementary Data files: 2 to 10, attached as separate files

**Supplementary Figures**

**Supplementary Figures S1a-d: sequence motif identification**
**Background:**
Two gene sets were selected for the motif searching: (Set 1) The top scoring 39 genes most highly correlated (Spearman) with the expression profile of the three cold-shock inducible transcriptional regulators (TFs) identified in our study: SCO1568, SCO4640 and SCO7014. Eleven cold-shock induced genes associated with membrane remodelling and efflux pumps plus the cold-shock inducible TF, SCO7014, were added to the list forming a total of 50 genes (Supplementary Data File S6 Genes correlating with the three CS TFs) ; (Set 2) Genes that were reported in the four functional enrichment categories identified in this study as putative members of the cold shock regulon (55 genes; Figures 4 and 5; Supplementary Data File S5).

Four conserved motifs and one conserved large inverted repeat which have potential regulatory roles in the cold shock response were identified (Figure S1a). The three major coldshock operons contain one or more motifs in the upstream region of genes encoding the CspA homologue and the DEAD box helicases (Figure S1b, Supplementary Table S1). *SCO4685* and *SCO5920* have several conserved motifs in their upstream regions: CS_IR_sco, a large stem loop IR sequence; additionally, CS_Motif_1_sco and CS_Motif_2_sco motifs are present in the translation initiation regions of *SCO3732*, *SCO4685* and *SCO5920*. Moreover CS_Motif_4_sco is present in the translation initiation region of *SCO4685* and *SCO5921*.

Interestingly, CS_Motif_1_sco overlaps the start codons of the respective genes (Supplementary Figure S1b). The relative positions of the identified motifs 1-4 suggests a role in translational regulation. The motif alignments of the four conserved motifs are presented in **Supplementary Table S1**; the motif logos are presented in Supplementary Figure S1a.

A fifth motif, CS_Motif_3_sco, has been mapped in the intergenic region of the phenylacetic acid degradation-encoding (PAA) operon between the two divergently transcribed genes, *SCO7470* and *SCO7471* (Supplementary Figure S1c). The PAA operon is co-ordinately induced by coldshock. This suggests an important role in stress resistance and cold adaptation, for example through the provision of TCA cycle intermediates.

**Legends to Supplementary figures S1a-d.**

CS_Motif_1_sco: 3 sites

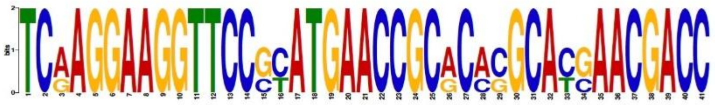

CS_Motif_2_sco: 28 sites

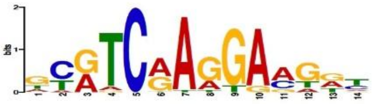

CS_Motif_3_sco: 7 sites

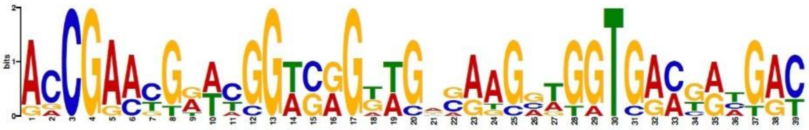

CS_Motif_4_sco: 8 sites

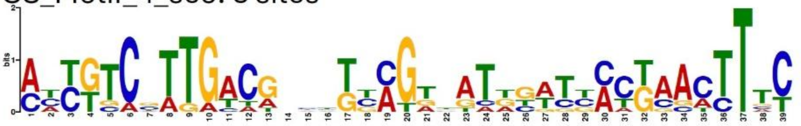

CS_IR_sco: 2 sites

```
                  ----------------- ->    <- -----------------
SCO5920 5'-TGTAGCTGGGGCCCGCATCCTTCGGGGTGCGGGCCCCAGCTACA
SCO4685 5'-TGCAGCTGGGGCCCGCATCCTTCGGGGTGCGGGCCCCAGCTGCA
```

**Figure S1a.** Conserved DNA sequence motifs identified in this study. Consensus logos of the cold-shock motifs 1 – 4 and the inverted repeat sequences upstream from the genes encoding the two major cold-shock induced helicases; blue font indicates nucleotide differences between the 2 CS_IR_sco sequences and red font indicates non-palindromic nucleotides.

#### Motifs identified upstream from cold-shock induced helicase-encoding genes

***SCO3732*. TSS upstream of *SCO3731* (2 TSS coordinates 4105236 and 4105198 upstream of *SCO3731*)**

CCGCCCCGCGGGGGCCACGGATTTCCCCCCGCGCATTCGACTTACCCGCGTTCCCCGCGT
TCCCCCGCATTCACCAATTCTCCCGGTCCGCTCGGCCCCACACGCCGAAATCGACCGGTC
CGGCTCGTTCTTGCGATTCCCTGCGCCGCTCGTCCGCTGCGGGTCTCCCTTGTTACGTGC
C*GTA**TCAAGGAAGGT***TCCCCATGAACCGCGCACGCACCAACGACCGCCGGCGCGCCGGCG
ACGGCCCCCGTCGCTCCCGTTCGGCCGGCCGCCCCCAGAACTCCGGCCGCCGCCCGGCCG

***SCO4685*, encoding DEAD box helicase.** TSS is upstream of ***SCO4684*** coordinate 5113589

------------------ -> <- -
5’-CCTGACGTTGGCGCGTCATCCGGCGCACACTT**TGCAGCTGGGGCCCGCATCCTTCGGGGT**
3’-GGACTGCAACCGCGCAGTAGGCCGCGTGTGAA**ACGTCGACCCCGGGCGTAGGAAGCCCCA**
-----------------
**5’-GCGGGCCCCAGCTGCA**TTCCCGTTCATCCGGCATTTCTCTTGCAATTCTCCACGTGCTCC
3’-**CGCCCGGGGTCGACGT**AAGGGCAAGTAGGCCGTAAAGAGAACGTTAAGAGGTGCACGAGG

5’-GTGCTCCGTGCTCCGTGCATCGTGCTCCGTGCTGCGG*GAATTCCTTGATAT*GTGCCGCAT
3’-CACGAGGCACGAGGCACGTAGCACGAGGCACGACGC**CCTTAAGGAACTATACACGGCGTA**

5’-T*GCG**TCAAGGAAGGT***TCCGCATGAACCGCACCGGCATGAACGACCGCATGAACGACCGCC
3’-**ACGCAGTTCCTTCCA**AGGCGTACTTGGCGTGGCCGTACTTGCTGGCGTACTTGCTGGCGG

5’-CCGCCCGCACGGGAAAGGCCCGCACGCGAGCCCTCGCGGTACAGGGTGAATTCGCCCACC
3’-GGCGGGCGTGCCCTTTCCGGGCGTGCGCTCGGGAGCGCCATGTCCCACTTAAGCGGGTGG

**>SCO5920 with 400 flanks (6487372 to 6489668)**
TSS upstream from ***SCO5921*** coordinate 6489982 73 nucleotides upstream from the start codon
**Stem-loop in black bold**

5’-CATCGCGCAGGGCCAGAAGGGCCCGACGGCCGAGAACATCGTTCCCGCCTGACGCCGACT
3’-GTAGCGCGTCCCGGTCTTCCCGGGCTGCCGGCTCTTGTAGCAAGGGCGGACTGCGGCTGA
------------------ -> <- ------------------
5’-GTCACGTACT**TGTAGCTGGGGCCCGCATCCTTCGGGGTGCGGGCCCCAGCTACA**TGCGTT
3’-CAGTGCATGA**ACATCGACCCCGGGCGTAGGAAGCCCCACGCCCGGGGTCGATGT**ACGCAA

5’-TCCCGCAGTGATTTCACCTGCGGACGACGCTCCGGACAGCGTGTGTTTACCGCACACGGA
3’-AGGGCGTCACTAAAGTGGACGCCTGCTGCGAGGCCTGTCGCACACAAATGGCGTGTGCCT

5’-TCGTCCTTCGAAGGTGCAGCCGCATCCTTCATAGCTACAGTGCAGTGGACGCTTCGAATA
3’-AGCAGGAAGCTTCCACGTCGGCGTAGGAAGTATCGATGTCACGTCACCTGCGAAGCTTAT

5’-TCGAGCCCACAGCACCCCCAATCATTCCAGCGCACGTGCCGCACAGATTTTCTGTCGGCC
3’-AGCTCGGGTGTCGTGGGGGTTAGTAAGGTCGCGTGCACGGCGTGTCTAAAAGACAGCCGG

5’-AGATCCGCTGGCAAGGCCCACCGGTCCATTCTTGCGATTCTCCGTGCTGCTCATTGCTTC
3’-TCTAGGCGACCGTTCCGGGTGGCCAGGTAAGAACGCTAAGAGGCACGACGAGTAACGAAG

5'-GG*AAATTCCTTGATAT*GTGCC*GCA****TCGAGGAAGGT****T**CCGTATGAACCGCACACGCACGAA**
3'-CC*TTTAAGGAACTATA*CACGG*CGTAGCTCCTTCCA*AGGCATACTTGGCGTGTGCGTGCTT

**5'-CGACC**GCTTCGCTCGCACCCGTCATGGCGGTGCCGACTCCGGAAAGGGCGGCAGCCGCTT
3'-GCTGGCGAAGCGAGCGTGGGCAGTACCGCCACGGCTGAGGCCTTTCCCGCCGTCGGCGAA

5'-CGGTTCGCCGGCGCCGCGCCGGCCGGCCGGACCCAGCCGCTCCGGCGGTTACGGCCGCCG
3'-GCCAAGCGGCCGCGGCGCGGCCGGCCGGCCTGGGTCGGCGAGGCCGCCAATGCCGGCGGC

**Figure S1b.** Conserved DNA sequence motifs identified upstream from cold-shock induced helicase-encoding genes. The different motifs listed in Table 1 are colour coded in the DNA sequence: **CS_Motif_1_sco** (3 sites, specific to the 3 helicase-encoding genes); **CS_Motif_2_sco**: this is a truncated version of **CS_Motif_1_sco** and is underlined and *italicised* in the sequence; **CS_Motif_4_sco**; **CS_IR_sco** shows the stem-loop sequence in *SCO4685* and *SCO5920*. Translation start codons, **ATG**, are indicated in **red.**

```
                                                          ← Sco7470
5′-CTGCTCAGTGGCCTGCGACACCGTATCCACCACCTTGTCTCCCGACCGACCATTCGGTTAGC
3′-GACGAGTCACCGGACGCTGTGGCATAGGTGGTGGAACAGAGGGCTGGCTGGTAAGCCAATCG

5′-ACGCGGCAAGGTCAAGTAATCCAGCCACGCGACGGGACTGTCAAGAGTGTGGCCGGCGAGGT
   TGCGCCGTTCCAGTTCATTAGGTCGGTGCGCTGCCCTGACAGTTCTCACACCGGCCGCTCCA
         < TSS
                                                          TSS >
5′-CCGGCTCTTGCCCGGCGCACGGAAACACGTCACACTGATACCGAACGAATGGTCGGTTGGGA
   GGCCGAGAACGGGCCGCGTGCCTTTGTGCAGTGTGACTATGGCTTGCTTACCAGCCAACCCT

                         → sco7471
5′-AGCTGGTGACGATGACCACGACACACGCCGCCGAGGCGCCGCCCCCGGACGCGCCGGACGGG
   TCGACCACTGCTACTGGTGCTGTGTGCGGCGGCTCCGCGGCGGGGGCCTGCGCGGCCTGCCC

5′-CTCCAGGAGCACTTCGACGCGACGATCGCGCGCGACCAGCGCATCGAGCCGCG-3′
   GAGGTCCTCGTGAAGCTGCGCTGCTAGCGCGCGCTGGTCGCGTAGCTCGGCGC-5′
```

**Supplementary Figure S1c.** Location of **CS_Motif_3_sco** DNA sequence motifs in the divergently transcribed *SCO7470-SCO7471*intergenic region of the phenylacetic acid degradation (PAA) regulon. Protein coding sequences are presented in boldface, TSSs (>) and start codons are boldface and underlined.

**CS_Motif_4_sco: 8 sites**

>**SCO3366** with 200 flanks (3723644 to 3725639) no TSS mapped
GCGGGTGTCCCCGCGGCGTCGCTGCTGCTTGGTGCCGTTCATCGTGCCGCCCATCCTCCT
ACGCACCTCCTGCCCCGCGCACTTTCCGAAAAACTT**ACTTGACGCCCGGCTAGTTACGCG**
**CCTACCTTTC**CCAGTGTAGACAACTTGCCGGGCGGCAAGTAAGTGGCAGTGGCACAGCGG
TAGCTGGGGGAGTAGGAGAG**ATG**GCCGAGAAAACGGAGGCGGCGACCGGTGGAGCCGGCG
CCGGCAAGCAGCCGAGGAGCGTGCGGGTCGTCCTGCTCGCCCTCATGATCGCGATGATGC

>**SCO3608** with 200 flanks (3985041 to 3985950) prx bidir-IG
CGGAACGCTCCCCGTATGAGTCCCCATTTGGAGAAGGACAGTACAGGCAACGGGTGCGGT
TCCAGGAGCGTCCGAAGAGATCGAGGGCGGCATT**AAAGGCGCATAAAGATTGCCGGAAGT**
TSS ✇
**CGGCAATGACGTG**CTGTCGCGCCGCGTGTGACGATTCGTTTC**A**CCGAAGCTCAGCGACGG
GCGGGAGGGGGACGGCACGC**ATG**GCCTGGTTTCTCGGGCTGGGTATCACGGGGGTCGTGC
TGCTCGCCCTGTCGCTGGTCTTCGACGGACTGCTGGAAGGCGTCTTCGACGGAGTGCTGG

>**SCO4121** with 200 flanks (4527845 to 4529522) no TSS mapped
AGGCGGCGATCTGCTCTTCGAGTGTCGGCTCGCCGGCGCCGGAGGTGTCGCCCATGGCCG
CAGTATGGCACGCAACTC**CTTTGCTTTGAAGTCCTTCGATATGTACTCTTCACCTTC**GAA
CTTTAGCTTCTAAGTCTTCAGCTCTAACGGCTGGAGCGGACGATCCGAGAGATCAAGAGA
GGGTGAACGTGACCAGGGCA**ATG**GGCGCGGCGATGCGCCGGATCCACGTGGGCAACGCAC
TCAGCGCGTTCGGACTCGGCTTCACGGTCCCCTACCTGTACGTCTATGTGGCGCAGGTAC

>**SCO4640** with 200 flanks (5064205 to 5065198) tetR - a top CS induced
CTCACGCTCTCCCTCGTCGGCGGGGGTCATCTCCTGTTCGCGGACCGCGACCCGGGCCAC
CCGCCGGCCGCGGCGGCCGTCGGCAAGCTGGTCGACACGGTGCTCGCCGGGGTCGTGCGG
CGACACCCGCGCTGACGCGGCACCGCCCTCCGCGCAC**ATCGTCCTTGACGAAGTTAGTGA**
TSS ✇ **leaderless**
**TTGAGTACTAACTTAC**TTTC**ATG**GTCAGGATGAGTGCAGAGGAGCGGCGCGAGAGCGTCA
TCCGTGCGGCGACGAGCGAGTTCGCCCGGGGTGGCTATCACGGCACGTCCACGGAGACCA

>**SCO4641** with 200 flanks (5064914 to 5066786) no TSS mapped
CCACCGCGTCTACGGCGGCATCTACGAGTCGGGACAGGCCAAGCCGAGCCGGAAGGACGC
GGACGCGCAGGACGCGGACGCCTAGGAACGAGCGCGGCACCCGCATACGGAACGACGACC
GCTTCCGCGCGCCCAC**AAAAGTTAGTCATCAATTACTAATCAAATCCACGGCAAG**TGCAT
GCACACCCTCTGGGGGAGAG**ATG**TCACAGCAAGGAGCACGTCGGAACACCCGCGGTGGGG
GAGCCGCCTGGGCCCTCGTCATCACCAGCGTCGCCGGCTTCATGGCGGCCCTGGACAACC

>**SCO4684** with 200 flanks (5113548 to 5114151) prx
TSS ✇
TCGGCGCGCAAAGACCGTAGCGGCGTGCT**ACTGTCGATATCAGTTGCAGTTGTGGTTCCC**
**GAATTTGC**AGCCCTTCCCGGCCGGTATTCGTGCCGTCGGAAGCGCTCTTTGACACCGGGT
TTTCCGGCGGGGTGATCATCACGGCGACACGGCGTTCGTACAGTGCGGACGCCGGCGCAC
TGCCCCAAAGGAGACAAGAC**ATG**GCTACTGGCACCGTGAAGTGGTTCAACGCGGAAAAGG
GATTCGGCTTCATCGAGCAGGACGGTGGCGGCGCGGACGTGTTCGCCCACTACTCGAACA

>**SCO4685** with 200 flanks (5113947 to 5115768) DEAD box helicase
CCTGACGTTGGCGCGTCATCCGGCGCACACTTTGCAGCTGGGGCCCGCATCCTTCGGGGT
GCGGGCCCCAGCTGCATTCCCGTTCATCCGGCATTTCTCTTGCAATTCTCCACGTGCTCC
GTGCTCCGTGCTCCGTGCATCGTGCTCCGTGCTGCG**GGAATTCCTTGATATGTGCCGCAT**
**TGCGTCAAGGAAGGT**TCCGC**ATG**AACCGCACCGGCATGAACGACCGCATGAACGACCGCC
CCGCCCGCACGGGAAAGGCCCGCACGCGAGCCCTCGCGGTACAGGGTGAATTCGCCCACC

>**SCO5921** with 200 flanks (6489417 to 6490020) prx

CCGCACAGTGGCTGCTGTGGCGTGCT**ACTGTCGATGTCAGTTGCAGTTGTGGTTCCCGAA**
**ATTTC**AAGTGCTCCGGTCGGGCTCAGTACCCCTGGAAGCACTTTTACATCTCCGGTCTTT

TTCCGGGCGGGGTGATCA**T**TGCGGCGACACGAGGTCCGCGCGGTGCGGATCCCGTACGAC
TGCCCCGAAGGAGAAATGAC**ATG**GCTGCTGGTACCGTGAAGTGGTTCAACGCGGAAAAGG
GCTTCGGCTTCATCGAGCAGGACGGTGGCGGCGCTGATGTGTTCGCCCACTACTCGAACA

**Supplementary Figure S1d.** CS_Motif_4_sco motif in cold-shock induced genes (8 sites).

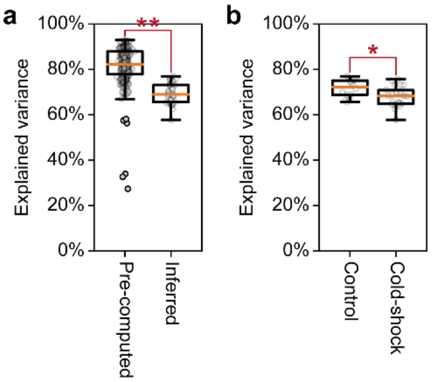


**Supplementary Figure S2. The percentage of transcriptomic variance explained by previously computed iModulons.** (a) Comparison of iModulon-explained transcriptomic variance between PRECISE-454 samples and cold-shock RNA-seq samples. (b) Comparison of iModulon-explained transcriptomic variance between cold-shock conditions and control conditions without cold-shock stress.

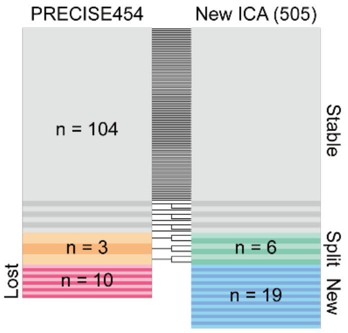


**Supplementary Figures S3. Comparison between previously computed iModulons and newly computed iModulons with cold-shock conditions.** To identify newly emerged or lost modules from the integration of cold shock conditions, cosine similarity between old and new iModulons was calculated; iModulon pairs with cosine similarity > 0.2 were linked together, indicating that the old iModulon has been retained, and the unlinked iModulons were regarded as lost or newly emerged modules. If multiple iModulons showed cosine similarity > 0.2 to a single original iModulon, they were considered as split modules.

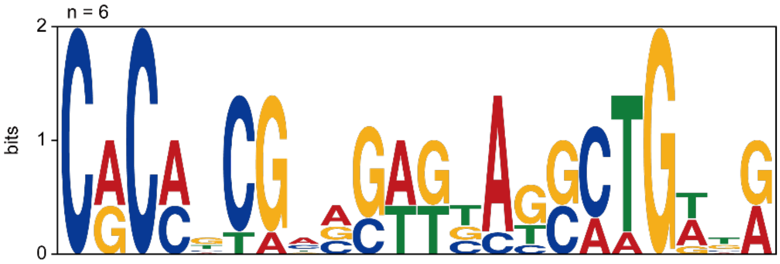


**Supplementary Figures S4. Sequence motif identified in the promoter regions of cold-shock regulatory iModulon genes.** Motif search was performed using the 50-nt upstream sequences from the transcription start sites reported by Jeong et al. with the MEME Suite using the ZOOPS option.

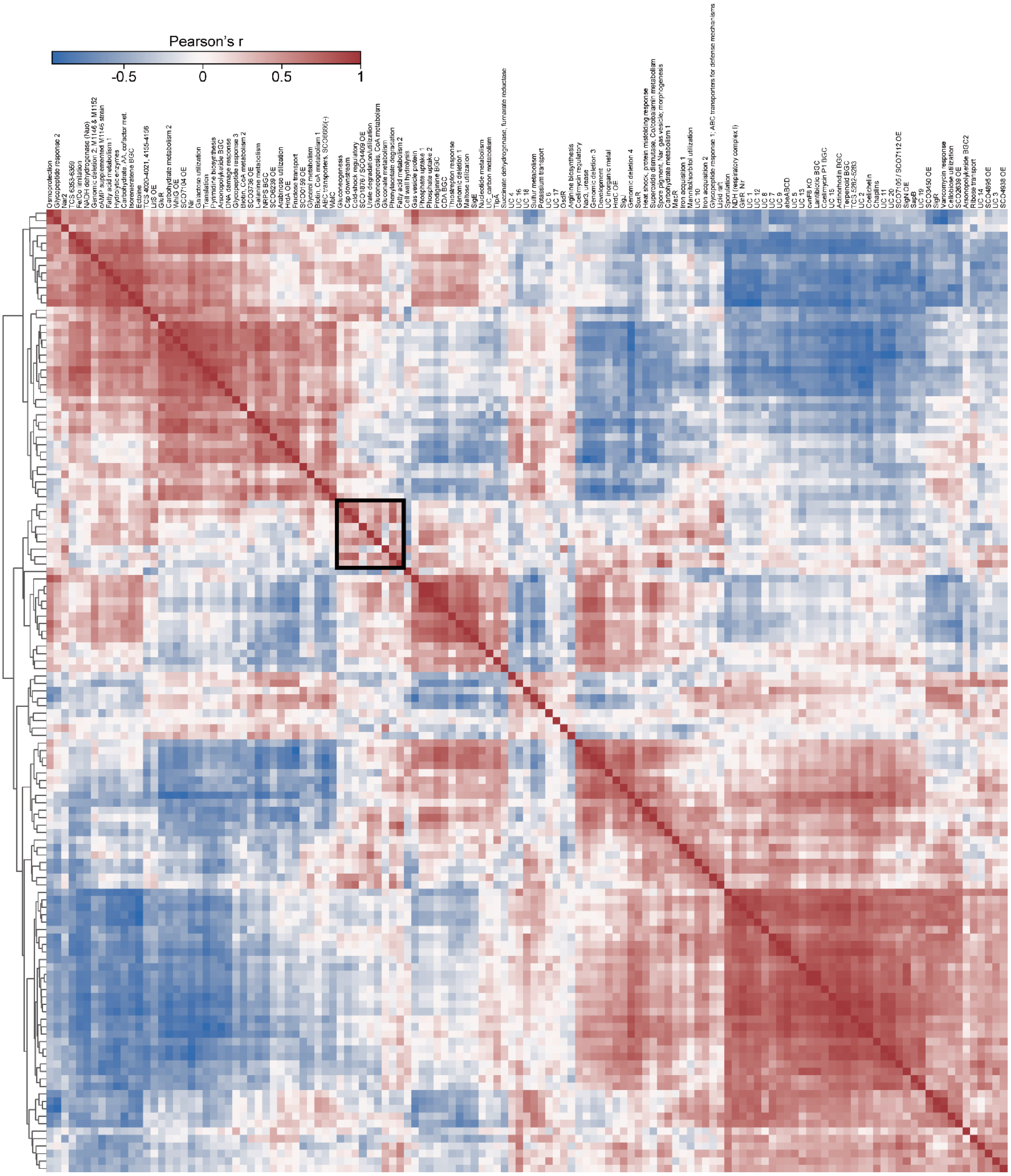


**Supplementary Figures S5. Hierarchical clustering of iModulon activities across cold-shock samples.** The black square indicates the subcluster containing the two cold-shock iModulons.

**a**

KEGG Pathways

| pathway | description | count in network | strength | signal | false discovery rate |
|---|---|---|---|---|---|
| sco00190 | Oxidative phosphorylation | 30 of 65 | 0.52 | 0.84 | 1.68e-05 |
| sco01100 | Metabolic pathways | 230 of 1021 | 0.21 | 0.81 | 3.17e-10 |
| sco01110 | Biosynthesis of secondary metabolites | 116 of 484 | 0.24 | 0.66 | 6.19e-06 |
| sco01200 | Carbon metabolism | 47 of 158 | 0.33 | 0.59 | 0.00029 |
| sco01230 | Biosynthesis of amino acids | 44 of 151 | 0.33 | 0.55 | 0.00067 |

**b**

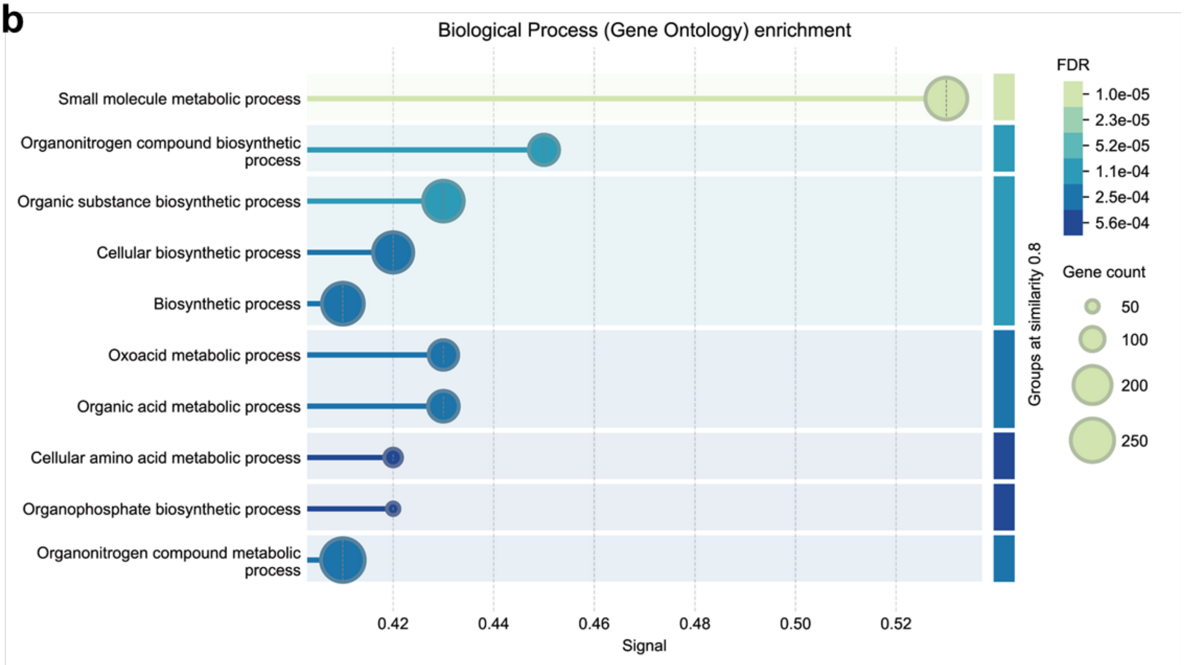


**Supplementary Figure S6. Pathways downregulated following cold-shock.** (a) KEGG pathways; (b) Biological Process (Gene Ontology) enrichment.

#### Supplementary Tables

#### Supplementary Table 1. Conserved DNA sequence motifs in cold-shock induced genes

**CS_Motif_1_sco: 3 sites**

Motif **TCRAGGAAGGTTCCSCATGAACCGCRCMSGCACSAACGACC** MEME-3 sites sorted by position p-value

| Sequence name | Strand | Start | P-value | | Site | |
|---|---|---|---|---|---|---|
| SCO5920 | + | 185 | 2.93e-24 | ATGTGCCGCA | TCGAGGAAGGTTCCGTATGAACCGCACACGCACGAACGACC | GCTTCGCTCG |
| SCO4685 | + | 185 | 7.05e-24 | CCGCATTGCG | TCAAGGAAGGTTCCGCATGAACCGCACCGGCATGAACGACC | GCATGAACGA |
| SCO3732 | + | 185 | 1.76e-23 | ACGTGCCGTA | TCAAGGAAGGTTCCCCATGAACCGCGCACGCACCAACGACC | GCCGGCGCGC |

**CS_Motif_2_sco: 28 sites**

Motif **GYRTCRAGGAMGDY** MEME-2 sites sorted by position p-value

| Sequence name | Strand | Start | P-value | | Site | |
|---|---|---|---|---|---|---|
| SCO3732 | + | 182 | 5.78e-09 | GTTACGTGCC | GTATCAAGGAAGGT | TCCCCATGAA |
| SCO5920 | - | 163 | 1.36e-08 | GATGCGGCAC | ATATCAAGGAATTT | CCGAAGCAAT |
| SCO4685 | + | 182 | 4.08e-08 | GTGCCGCATT | GCGTCAAGGAAGGT | TCCGCATGAA |
| SCO5920 | + | 182 | 1.03e-07 | GATATGTGCC | GCATCGAGGAAGGT | TCCGTATGAA |
| SCO4640 | - | 158 | 1.80e-07 | ATCACTAACT | TCGTCAAGGACGAT | GTGCGCGGAG |
| SCO4685 | - | 158 | 2.01e-07 | AATGCGGCAC | ATATCAAGGAATTC | CCGCAGCACG |
| SCO3366 | - | 92 | 2.31e-07 | AACTAGCCGG | GCGTCAAGTAAGTT | TTTCGGAAAG |
| SCO0800 | + | 189 | 3.65e-07 | TGTTGAACCC | GCGTCAAGTAAGGT | GCGGGGCATG |
| SCO7170 | - | 10 | 1.45e-06 | GCGGACCGCG | GTGACAAGGAAGGT | GGAGGCCGC |
| SCO2737 | - | 12 | 1.45e-06 | GGCCTTCAGG | GCGTCGAGGAAGGA | GGTGAAAGCC |
| SCO0800 | - | 93 | 1.45e-06 | ACACGCCGCT | GCGTCAAGGGAGAT | CGCGTCATGG |
| SCO7533 | + | 217 | 2.43e-06 | CGAGTCCTGC | TCATCGAAGACGAC | CCCGCGGTGC |
| SCO7230 | + | 217 | 2.43e-06 | CGGGTCCTGA | TCATCGAAGACGAC | CGCGCCGTCC |
| SCO3471 | + | 179 | 3.88e-06 | AGCCAGGCGG | GAATCGAAGAAGGA | GAACGATCGT |
| SCO7794 | - | 246 | 4.74e-06 | CATGCCCTGG | TCGTCAAGGAGTTC | GAGCGCCGCC |
| SCO3682 | + | 258 | 6.16e-06 | CTTCGCCCGG | CTGTCGAAGAAGGT | CGCGGATGCC |
| SCO3956 | - | 183 | 6.63e-06 | GATCACAGTC | ACGTCAATGACAGT | AGTGACGTCC |
| SCO5912 | - | 15 | 8.44e-06 | ACAGGGAATG | GCATCGAGGAGTGA | GACCGTGGGC |
| SCO7475 | + | 185 | 9.18e-06 | CCGTTCGAGT | ACGTCAAGGAGATC | TGATGGCCCC |
| SCO5517 | + | 147 | 9.80e-06 | TCTTATACGC | GTGTCTAGGACATT | GTCTAAGACG |
| SCO4641 | + | 146 | 1.05e-05 | CAAAAGTTAG | TCATCAATTACTAA | TCAAATCCAC |

| | | | | | | |
|---|---|---|---|---|---|---|
| SCO7472 | + | 62 | 1.05e-05 | CTGATGCGCG | TCATCAAGGGCGAC | GGCCCCTGCA |
| SCO5862 | + | 214 | 1.14e-05 | CGCGTACTCG | TCGTCGAGGACGAG | CAGCTGCTCG |
| SCO7473 | - | 153 | 1.22e-05 | GTCGTCGGGG | ATGTCGTAGAAGGT | CGGGTGGCGG |
| SCO3685 | + | 79 | 1.64e-05 | TTGTCCGGGC | GCATCGAGGAGTAG | CCTGAGACGT |
| SCO7014 | + | 222 | 1.64e-05 | ACTTGCTCAG | GTGGCGAAGAAGGT | TGGGGTCAGC |
| SCO3732 | - | 163 | 2.16e-05 | GATACGGCAC | GTAACAAGGGAGAC | CCGCAGCGGA |
| SCO6262 | + | 153 | 2.78e-05 | CCCGCCCGTA | GAGTCGAAGACATC | TGATCGGGTG |

--------------------------------------------------------------------------------

**CS_Motif_3_sco: 7 sites**

Motif **ACCGAACGRWCGGWSRGKWGRSAAGVWGGTGAMGRHGAC** MEME-1 sites sorted by position p-value

| Sequence name | Strand | Start | P-value | | Site | |
|---|---|---|---|---|---|---|
| SCO7471 | + | 167 | 3.67e-20 | TCACACTGAT | ACCGAACGAATGGTCGGTTGGGAAGCTGGTGACGATGAC | CACGACACAC |
| SCO7470 | - | 32 | 3.67e-20 | TCACACTGAT | ACCGAACGAATGGTCGGTTGGGAAGCTGGTGACGATGAC | CACGACACAC |
| SCO7471 | - | 23 | 1.46e-16 | CCGCGTGCTA | ACCGAATGGTCGGTCGGGAGACAAGGTGGTGGATACGGT | GTCGCAGGCC |
| SCO7470 | + | 176 | 1.46e-16 | CCGCGTGCTA | ACCGAATGGTCGGTCGGGAGACAAGGTGGTGGATACGGT | GTCGCAGGCC |
| SCO3167 | + | 173 | 5.12e-13 | GAGTGGACCG | GCCGACCGGACGGAGAGGAGTGAGCGAGATGACGGAGAC | CGCCACCGCA |
| SCO4121 | + | 155 | 5.70e-13 | TAACGGCTGG | AGCGGACGATCCGAGAGATCAAGAGAGGGTGAACGTGAC | CAGGGCAATG |
| SCO4871 | - | 112 | 2.41e-12 | GACGGGACTT | AACGACGTTAAGGAGAGTTGCCGTGAATGTCACGGATAC | GCATGTCGCG |

**CS_Motif_4_sco: 8 sites**

Motif **MHYKTCSTTGACGNHHKBMGDBATDSWTBMCKAACTTTC** MEME-2 sites sorted by position p-value

| Sequence name | Strand | Start | P-value | | Site | |
|---|---|---|---|---|---|---|
| SCO4640 | + | 158 | 2.21e-17 | CTCCGCGCAC | ATCGTCCTTGACGAAGTTAGTGATTGAGTACTAACTTAC | TTTCATGGTC |
| SCO5921 | + | 27 | 2.58e-15 | GTGGCGTGCT | ACTGTCGATGTCAGTTGCAGTTGTGGTTCCCGAAATTTC | AAGTGCTCCG |
| SCO4641 | - | 137 | 2.51e-14 | TGTGCATGCA | CTTGCCGTGGATTTGATTAGTAATTGATGACTAACTTTT | GTGGGCGCGC |
| SCO4121 | + | 79 | 2.74e-13 | CACGCAACTC | CTTTGCTTTGAAGTCCTTCGATATGTACTCTTCACCTTC | GAACTTTAGC |
| SCO4684 | + | 30 | 3.58e-13 | GCGGCGTGCT | ACTGTCGATATCAGTTGCAGTTGTGGTTCCCGAATTTGC | AGCCCTTCCC |
| SCO4685 | - | 157 | 2.81e-12 | TTCATGCGGA | ACCTTCCTTGACGCAATGCGGCACATATCAAGGAATTCC | CGCAGCACGG |
| SCO3608 | - | 95 | 2.90e-11 | GGCGCGACAG | CACGTCATTGCCGACTTCCGGCAATCTTTATGCGCCTTT | AATGCCGCCC |
| SCO3366 | + | 92 | 1.05e-10 | CTTTCCGAAA | AACTTACTTGACGCCCGGCTAGTTACGCGCCTACCTTTC | CCAGTGTAGA |

--------------------------------------------------------------------------------

**Supplementary Table S2. Gene membership of the 'cold-shock regulatory' iModulon.**

| Locus tag | Function | Weight |
|---|---|---|
| SCO5921 | cold-shock domain-containing protein | 0.252976 |
| SCO4684 | cold shock protein | 0.227367 |
| SCO4685 | DEAD/DEAH box helicase | 0.191994 |
| SCO1568 | TetR family transcriptional regulator | 0.166492 |
| SCO5920 | DEAD/DEAH box helicase | 0.141639 |
| SCO4908 | RNA polymerase sigma factor | 0.138803 |
| SCO4640 | TetR family transcriptional regulator | 0.125248 |
| SCO3683 | hypothetical protein | 0.118657 |
| SCO3684 | hypothetical protein | 0.103977 |
| SCO5746 | hypothetical protein | 0.082299 |
| SCO3685 | hypothetical protein | 0.08181 |
| SCO4504 | methyltransferase | 0.080563 |
| SCO1567 | transmembrane-transport protein | 0.079445 |
| SCO1145 | MarR family regulatory protein | 0.078669 |
| SCO0593 | hypothetical protein | 0.078338 |
| SCO4189 | hypothetical protein | 0.076768 |
| SCO4641 | transmembrane efflux protein | 0.07646 |
| SCO3681 | hypothetical protein | 0.076209 |
| SCO4054 | hypothetical protein | 0.075684 |
| SCO3732 | DEAD/DEAH box helicase | 0.074654 |
| SCO5987 | hypothetical protein | 0.0735 |
| SCO5484 | small hydrophobic membrane protein | 0.069553 |
| SCO3086 | lipoprotein | 0.067474 |
| SCO4107 | hypothetical protein | 0.065126 |
| SCO7014 | LacI family transcriptional regulator | 0.064682 |

**Supplementary Data File S1**: Oligonucleotides used for depletion of rRNAs and selected non-coding RNAs.

All probes listed below are given 5’-to-3’ and remove rRNAs and *rnpB, ssrA* and *srp* RNAs. Note: tRNAs are not removed by these probes.

| |
|---|
| AAGGAGGTGATCCAGCCGCACCTTCCGGTACGGCTACCTTGTTACGACTTCGTC |
| AAGTCGTTACGCCATTCGTGCAGGTCGGAACTTACCCGACAAGGAATTTCGCTA |
| AAGTGGTTCATCGTTCGACTTGCATGTGTTAAGCACGCCGCCAGCGTTCGTCCTGA |
| AATAATCCGACCGTTCACAGCGTCCTCGCTGTTGTGTTACTTCAAAGGAA |
| AATCACTAGACCAGTGAGCTATTACGCACTCTTTCAAGGGTGGCTGCTTCTAAGC |
| AATCAGGGCTTCTCCGTGTGCAGTCCGCTTCGATTTTCTCGGCCCCGGAGATCACG |
| AATGCTGATCTGCGATTACTAGCGACTCCGACTTCATGGGGTCGAGTTGCAGAC |
| ACAACGTGGAATGTTGCCCACACCTAGTGCCCACCGTTTACGGCGTGGACTACCA |
| ACAGCCATGCACCACCTGTACACCGACCACAAGGGGGGCACCATCTCTGATGCTTTCC |
| ACATAAGGGGCATGATGACTTGACGTCGTCCCCACCTTCCTCCGAGTTGACCCCG |
| ACCTCGCAACACACCGCAAACTCGCAGGCTCATTCTTCAAAAGGCACGCAGTCACGACG |
| ACGCTTTCGCTCCTCAGCGTCAGTATCGGCCCAGAGATCCGCCTTCGCCACCGGTGT |
| ACTACGGGGGTCTTACCCTCTACGCCGGACCTTTCGCATG |
| ACTCAACACCTGATTGCCAACCAGGCTGAGGGAACCTTTGGGCGCCTCCGTTACT |
| ACTTAATGCGTTAGCTGCGGCACGGACAACGTGGAATGTTGCCCACACCTAGTGCCCA |
| AGAACCAGCTATCACGGAGTTTGATTGGCCTTTCACCCCTAACCACAGGTCATC |
| AGACTGGTATTTCAACGACGACTCCACCCACACTGGCGTGCGAGCTTCAAAGTCT |
| AGGACAAGCCCTCGGCCTATTAGTACCGGTCACCTCCACA |
| ATAATTCCGGACAACGCTTGCGCCCTACGTATTACCGCGGCTGCTGGCACGTAGTTA |
| ATATCTGCGCATTTCACCGCTACACCAGGAATTCCGATCTCCCCTACCGAACTCTAG |
| ATCACTCCGCTTCGGGTCTTGAGCGTGCTACTAAAAACGC |
| ATCCACCGTGCGCCCTTAAAAACTTGGCCACAGATGCTCG |
| ATCCTTCACCGCCGGAGCTTTCGAACCTCGCAGATGCCTGCGAGGGTCAGTATCC |
| ATGCTCTGGGCTGTTTCCCTCTCGACCATGGAGCTTATCCCCCACAGTCTCACTG |
| ATGTAACCGGGCGTTTCCCCTACGCTATGACCACCGAAAC |
| ATTAAGCCACATGCTCCGCCGCTTGTGCGGGCCCCCGTCAATTCCTTTGAGTTTTA |

| |
|---|
| ATTACTAGCGACTCCGACTTCATGGGGTCGAGTTGCAGACCCCAATCCGAACTGAGA |
| ATTGTACCGGCCATTGTAGCACGTGTGCAGCCCAAGACATAAGGGGCATGATGACTTGA |
| ATTGTGCAATATTCCCCACTGCTGCCTCCCGTAGGAGTCTGGGCCGTGTCTCAGTC |
| CAAACCATCCCGTCGATATGGACTCTTGGGGAAGATCAGCCTGTTATC |
| CAAACGTTTCTCACGTTTGTATCGCTACTCATGCCTGCATTCTCACTC |
| CAACACCCCGAAGGGCTTGCTGGCAACACGGGACAAGGGTTG |
| CAACACGGGACAAGGGTTGCGCTCGTTGCGGGACTTAACC |
| CAAGAAACACACGACGCTGCACCTAAATGCATTTCGGGGAGAACCAGCTATCA |
| CAAGGCATCCACCGTGCGCCCTTAAAAACTTGGCCACAGA |
| CAAGGTGTGCGGATTTACCTACACACCGGCCTACACCCTTAC |
| CAAGTCCTTGCGCGACGCTCCCACGGCTTGTAGGCACACGGTTTCAGGTACTATTT |
| CAATATCAAACTGTAGTAAAGGTCCCGGGGTCTTTCCGTCCTGCTGCGCGAAA |
| CAATCCGAACTGAGACCGGCTTTTTGAGATTCGCTCCACCTTGCGGTAT |
| CACACCAGAGGTTCGTCCGTCCCGGTCCTCTCGTACTAGG |
| CACACGCAACCCCTGCCGGGTCTCACACGTATACGGTTTG |
| CACACTGGCGTGCGAGCTTCAAAGTCTCCCAGCTATCCTACACAAG |
| CACATCCTTTCCCACTTAGCGTACGCTTAGGGGCCTTAGTCGATGCTCTG |
| CACCCCCAGCTCAGAGTGCAAAACTCGTCACCAGGTGTGG |
| CACCCGGGTTAACCTCGCAACACACTGCAAACTCGCAGGC |
| CACCGAAGTGGTTCATCGTTCGACTTGCATGTGTTAAGCAC |
| CACCGTTTACGGCGTGGACTACCAGGGTATCTAATCCTGTTC |
| CACCTTCCTCCGAGTTGACCCCGGCGGTCTCCCGTGAGTC |
| CACCTTCGACAGCTCCCTCCCACAAGGGGTTGGGCCACCGGCTTCGGGTGTTA |
| CACGCACCCGCCAGAGCCGACCTACCCTTGCTGCCTTCCG |
| CACGCAGTCACGAGACACCAAGCAAGCTTGATGTCCGACGCTCCCACGGCTTGTAGGCAC |
| CACGCAGTCACGAGGATGGAGCAAGCTCCATCCCGACGCTC |
| CACGCAGTCACGAGGATGGAGCAAGCTCCATCCCGACGCTCCCACGGCTTGTAGGCA |
| CACGCTTTCGCTCCTCAGCGTCAGTATCGGCCCAGAGATC |
| CACTCCCCTCAACTCCGAAGAGATCAGGGCGGCTTCACGG |
| CAGACCCCAATCCGAACTGAGACCGGCTTTTTGAGATTCG |
| CAGACCGTAAAAGAGAGATCCCACAACCCCGCACACGCAACCCCTGCCGGGTCTCA |
| CAGACGAGTCGGGCTGTACGCCGGGTTCTGTTCCGACGGG |
| CAGAGATCCGCCTTCGCCACCGGTGTTCCTCCTGATATCTG |

| |
|---|
| CAGATGCTCGCGTCCACTGTGCAGTTCTCAAACAACGACC |
| CAGATTGCCCACGTGTTACTCACCCGTTCGCCACTAATCC |
| CAGCTATCCTACACAAGCCGAACCGAACACCAATATCAAACTGTAGTAAA |
| CAGCTCAGAGTGCAAAACTCGTCACCAGGTGTGGCCCCCCTTCTCCCGAAGTTA |
| CAGGAACCCTTAGTCAATCGGCGCAAACGTTTCTCACGTTT |
| CAGTACCATCGGCGCTGTAAGGCTTAGCTTCCGGGTTCGAAATGTAAC |
| CAGTCACGACGCAAGGACAAGTCCTTGCGCGACGCTCCCAC |
| CAGTCACGAGACACCAAGCAAGCTTGATGTCCGACGCTCC |
| CAGTCACTGTTTGGTTTCCCTCTTCACCCCGTGACCGGGATCAA |
| CAGTCGTCTACTGGGAGCCTTAACCCCTCAAGGGGGTGGGAGTCCTCATCTCGAA |
| CAGTCTCACTGCCGCGCTCTCACTTACCGGCATTCGGAGTTTG |
| CAGTTAAACTACCCATCAGACACTGTCCCTGATCCGGATCAC |
| CATCACAGCCGGCGTTGGCCGTATTGCTGCAATGACACGACTTC |
| CATCACTCACCTACTAACCGCTTGGTTCGGCGGCTCCACC |
| CATCAGGTCTCAGACTCATGTCAGGCGGATTTACCTACCTGAC |
| CATCAGGTCTCAGCCACAAGGTGTGCGGATTTACCTACAC |
| CATCCATCTAGGACCGGCGTTGCCACCGGCCTCCAGCGGTCTAC |
| CATCCTTCACCGCCGGAGCTTTCGAACCTCGCAGATGCCT |
| CATCGCCTATCCAGTGCTCTACCTCCGGCAAGAAACACAC |
| CATCTCTGATGCTTTCCGGTGTATGTCAAGCCTTGGTAAGGTTCTTC |
| CATGCCCTTGGCAGGACAACTGGCACACCAGAGGTTCGTC |
| CATTACCTCACCAACAAGCTGATAGGCCGCGGGCTCATCCTTC |
| CATTCGGACACCCCCGGATCAAAGCCTGGTTGACGACTCC |
| CATTCGGAGTTTGGCTAAGGTCAGTAACCCGGTAGGGCCCATCG |
| CATTCTTCAAAAGGCACGCAGTCACGAGGATGGAGCAAGC |
| CCAAGCAAGCTTGATGTCCGACGCTCCCACGGCTTGTAGGCACACGGTTTCAGGTACTA |
| CCACAATCGGCTCGGCATCAGGTCTCAGCCACAAGGTGTG |
| CCAGGATCAAACTCTCCGTGAATGTATACCCGTAATCGGG |
| CCAGGATGCGACGAGCCGACATCGAGGTGCCAAACCATCC |
| CCATCACTCACCTACTAACCGCTTGGTTCGGCGGCTCCAC |
| CCATGAAACTCGCTAGAGGCTTTTCTCGACAGCATAGGATCATC |
| CCGAAGCCTCCGGCGGTCTGCTTTCTGTGGCACTGTCCCG |
| CCGGCCTACACCCTTACCCCGGGACAACCACCGCCCGGGATGGACTACC |

| |
|---|
| CCGGGTTAACCTCGCAACACACCGCAAACTCGCAGGCTCATTCTTCAAAAGGCACGC |
| CCGTCAATTCCTTTGAGTTTTAGCCTTGCGGCCGTACTCC |
| CGCACCTTCCGGTACGGCTACCTTGTTACGACTTCGTCCCAATCGCCAGTCCCA |
| CGCTCGTTGCGGGACTTAACCCAACATCTCACGACACGAG |
| CGGAATCACTAGACCAGTGAGCTATTACGCACTCTTTCAAGG |
| CGGTGTTCCTCCTGATATCTGCGCATTTCACCGCTACACC |
| CGTATTGCTGCAATGACACGACTTCGGCGGTACGCTTGAG |
| CGTCCTACTCTCCCACAGGGTCCCCCTGCAGTACCATCG |
| CGTCGCTGCATCAGGCTTTCGCCCATTGTGCAATATTCCC |
| CTAACCACAGGTCATCCCCCAGGTTTTCAACCCTGGTGGGTTCGGTCC |
| CTACATTGTCGGCGCGGAATCACTAGACCAGTGAGCTATTA |
| CTACCGAACTCTAGCCTGCCCGTATCGACTGCAGACCCGGGGTTAAGCCCCG |
| CTACCTTGTTACGACTTCGTCCCAATCGCCAGTCCCACCTTCG |
| CTACCTTGTTACGACTTCGTCCCAATCGCCAGTCCCACCTTCGACAG |
| CTACCTTGTTACGACTTCGTCCCAATCGCCAGTCCCACCTTCGACAGCTCCCT |
| CTACGGCTTCCCCACCCGGGTTAACCTCGCAACACACCGC |
| CTACGGCTTCCCCACTCGGGTTAACCTCGCAACACACCGC |
| CTACGTATTACCGCGGCTGCTGGCACGTAGTTAGCCGGCGCTTCTTCTG |
| CTACTAACCGCTTGGTTCGGCGGCTCCACCACTCCCCTCAACTCCGAAGAGATCA |
| CTACTTGGGTGTCTCTCAAACGAGCCGTTGACGTTTCGACTAC |
| CTAGAGGCTTTTCTCGACAGCATAGGATCATCCACTTCACCACAATC |
| CTATCAACCCAGTCGTCTACTGGGAGCCTTAACCCCTCAAGG |
| CTATGGTTGAGACAGTCGAGAAGTCGTTACGCCATTCGTG |
| CTATTAGTACCGGTCACCTCCACACCTCACGGTGCTTCCAGATCCGGCCTATCAACC |
| CTCACGGTACTATCCGCTATCGGTCACCAGGGAATATTTAGGCTTAG |
| CTCATCTCGAAGCAGGCTTCCCGCTTAGATGCTTTCAGCGGTTATC |
| CTCCAGCGGTCTACCCGCGGACTCGGGCGGGCAGCCCTCGAACGTCCGCGCAGAAG |
| CTCCCACGTCCTTCATCGGTTCCTGGTGCCAAGGCATCCAC |
| CTCGAACGTCCGCGCAGAAGCACCGGGGTGCTTCCTTCTTGACCTTGCTC |
| CTCGGGTTAACCTCGCAACACACCGCAAACTCGCAGGCTCATTCTTCAAAAGGCACGC |
| CTCGGTATTCTCTACCTGACCACCTGAGTCGGTTTAGGGTAC |
| CTCTCACTTACCGGCATTCGGAGTTTGGCTAAGGTCAGTAAC |
| CTCTCAGGCCGGCTACCCGTCGTCGCCTTGGTGAGCCATTAC |

| |
|---|
| CTCTTACAGCCGCTTCAACCTGCCCATGGCTAGATCACTC |
| CTCTTTACGCCCAATAATTCCGGACAACGCTTGCGCCCTACGTATTAC |
| CTGATATCTGCGCATTTCACCGCTACACCAGGAATTCCGATCTC |
| CTGCGCGAAACGAGCATCTTTACTCGTAGTGCAATTTCAC |
| CTGCTGGGAGGAATCGCGCTTGGTGTTGGCGATTATTTTT |
| CTGGGCACTGGTGGTCTCTTACACCACCGTTTCACCCTTAC |
| CTGTAAGGCTTAGCTTCCGGGTTCGAAATGTAACCGGGCGTTTCCCCTAC |
| CTGTGGAGCCCGGACGTTCCTCGGGAAGTCCCGTGAGGGACTC |
| CTTAACCCAACATCTCACGACACGAGCTGACGACAGCCATG |
| CTTAACGTTCCAGCACCGGGCAGGCGTCAGTCCGTATACATC |
| CTTAATGCGTTAGCTGCGGCACGGACAACGTGGAATGTTG |
| CTTACCCTCTACGCCGGACCTTTCGCATGTCCTTCGCCTACATC |
| CTTAGATGCTTTCAGCGGTTATCCCTCCCGAACGTAGCCAAC |
| CTTAGGATGGTTATAGTTACCACCGCCGTTTACTGGCGCTTAAGTTCTCA |
| CTTAGGGGCCTTAGTCGATGCTCTGGGCTGTTTCCCTCTCG |
| CTTAGGTCCCGACTTACCCTGGGCAGATCAGCTTGACCCAG |
| CTTAGTAGGAGGCTCGCTTCGCTACCTGAGATCAGGCAGCGAGGGCGAAGG |
| CTTAGTCAATCGGCGCAAACGTTTCTCACGTTTGTATCGCTACTCATG |
| CTTAGTCGATGCTCTGGGCTGTTTCCCTCTCGACCATGGAGCTTATC |
| CTTCAAAGTCTCCCAGCTATCCTACACAAGCCGAACCGAACACCAATA |
| CTTCAACCTGCCCATGGCTAGATCACTCCGCTTCGGGTCTTGAG |
| CTTCCACAAGCCACCGCCGGATCACTAGTCCCGACTTTCGTCCCTG |
| CTTCCAGATCCGGCCTATCAACCCAGTCGTCTACTGGGAG |
| CTTCCCCTGCTTCGACAGCCGCTGTCGAAACCGATCATCC |
| CTTCTAAGCCAACCTCCTGGTTGTCTGTGCGACTCCACATCCTTTC |
| CTTCTCAAGACTCCTACGCGCACAGCGGATAGGGACCGAAC |
| CTTCTCCCGAAGTTACGGGGGCATTTTGCCGAGTTCCTTAACCA |
| CTTCTTCCCTGCTGAAAGAGGTTTACAACCCGAAGGCCGTCATC |
| CTTGAGCGTGCTACTAAAAGCGCCCTATTCGGACTCGCTT |
| CTTGTAGGCACACGGTTTCAGGTACTATTTCACTCCGCTCCC |
| CTTGTAGGCACACGGTTTCAGGTACTATTTCACTCCGCTCCCG |
| CTTGTCCCAGAGTGAAGGGCAGATTGCCCACGTGTTACTC |
| CTTGTGCACTTACACTCAACACCTGATTGCCAACCAGGCTGAGGGAACCTTT |

| |
|---|
| CTTTAATGGGCGAACAGCCCAACCCTTGGGACCGACTCCAG |
| CTTTCACAACCGACGTGACAAGCCGCCTACGAGCTCTTTACGCCCAATAATT |
| CTTTCGCATGTCCTTCGCCTACATCAACGGTTTCTGACTC |
| GAAACGAGCATCTTTACTCGTAGTGCAATTTCACCGGGCCTATGGTTGAG |
| GAAAGGAGGTGATCCAGCCGCACCTTCCGGTACGGCTACC |
| GAAATGTGACTAACCGGTCCCCTTAACGTTCCAGCACCGGGCAGGC |
| GAACAGCCCCGGGCTGACACTCCCTTAGTAGGAGGCTCGCTTC |
| GAACCGAACACCAATATCAAACTGTAGTAAAGGTCCCGGGGTCTTTC |
| GAACCTTTGGGCGCCTCCGTTACTCTTTAGGAGGCAACCG |
| GAACGTATTCACCGCAGCAATGCTGATCTGCGATTACTAG |
| GAACTGTCTCACGACGTTCTAAACCCAGCTCGCGTACCGCTTTAATG |
| GAAGAGATCAGGGCGGCTTCACGGCCTTAGCATCACGATG |
| GAATATCAACCGGTTATCCATCGACTACGCCTGTCGGCCTCGCCTTAGGTC |
| GAATATTTAGGCTTAGCGGGTGGTCCCGCCAGATTCACACGGGATTTCTC |
| GAATCACGGTTGTTTTCTCTTCCTGCGGGTACTGAGATGTTTCACTTC |
| GAATCGAACCCGCGTCCAACGGTGCGGAATCAGGGCTTCTC |
| GACAACCACCGCCCGGGATGGACTACCTTCCTGCGTCACCCCAT |
| GACAAGGAATTTCGCTACCTTAGGATGGTTATAGTTACCACCGCCGTTTA |
| GACATAAGGGGCATGATGACTTGACGTCGTCCCCACCTTC |
| GACATCGAGGTGCCAAACCATCCCGTCGATATGGACTCTTG |
| GACCCGTCGGTCTCACAGTCAAGCTCCCTTGTGCACTTAC |
| GACTACCAGGGTATCTAATCCTGTTCGCTCCCCACGCTTTCG |
| GACTTACCCTGGGCAGATCAGCTTGACCCAGGAACCCTTAGTCAATC |
| GACTTTCGTCCCTGCTCGACCCGTCGGTCTCACAGTCAAG |
| GAGATCACGCGGACAAGTCTCCGACGGGCCCAGTCACTGT |
| GAGCTTTCGAACCTCGCAGATGCCTGCGAGGGTCAGTATCCGGTATTAGACCCCGTT |
| GAGTTCCTTAACCATAGTTCACCCGAACGCCTCGGTATTCTCTACCTGAC |
| GAGTTTGGCTAAGGTCAGTAACCCGGTAGGGCCCATCGCCTATCCAGTGCTCTA |
| GATAGGCCGCGGGCTCATCCTTCACCGCCGGAGCTTTCGA |
| GATAGGGACCGAACTGTCTCACGACGTTCTAAACCCAGCTC |
| GATCAAACTCTCCGTGAATGTATACCCGTAATCGGGTGAC |
| GATCAAAGCCTGGTTGACGACTCCCCGGGGCCTATCGTGG |
| GATCAAGGTTTAGTCCCCTAGCTGATGCCAGGATCCGGGTC |

| |
|---|
| GATCACGGACCCAGGTTAGACATCCAGCACGACCAGACTG |
| GATCACTCCGCTTCGGGTCTTGAGCGTGCTACTAAAACGC |
| GATCTCCCCTACCGAACTCTAGCCTGCCCGTATCGACTGCAGAC |
| GATGGACTACCTTCCTGCGTCACCCCATCACTCACCTACTAAC |
| GATGGAGCAAGCTCCATCCCGACGCTCCCACGGCTTGTAGGCACACGGTTTCAGGTACT |
| GATTTCTCGGGCCCCGTGCTACTTGGGTGTCTCTCAAACGAG |
| GCAGATGCCTGCGAGGGTCAGTATCCGGTATTAGACCCCGTTTCCAGGGCTTGTCCCA |
| GCAGGCGTCAGTCCGTATACATCGCCTTACGGCTTCGCAC |
| GCCACCGGCTTCGGGTGTTACCGACTTTCGTGACGTGACG |
| GCCAGATTCACACGGGATTTCTCGGGCCCCGTGCTACTTG |
| GCCTATTAGTACCGGTCACCTCCACACCTCACGGTGCTTCCAGATCCGGCCTATCAAC |
| GCGGATTTACCTACCTGACGTCCTACACCCTTACCCCGGGACAACCACCGC |
| GCGTGCTACTAAAACGCCCTATTCGGACTCGCTTTCGCTACGGCTTCC |
| GCTCCCCTACCCATCACAGCCGGCGTTGGCCGTATTGCTG |
| GCTTATCCCCCACAGTCTCACTGCCGCGCTCTCACTTACC |
| GGACAAGCCCTCGGCCTATTAGTACCGGTCACCTCCACAC |
| GGAGAACCAGCTATCACGGAGTTTGATTGGCCTTTCACCC |
| GGCATTTTGCCGAGTTCCTTAACCATAGTTCACCCGAACG |
| GGCCTTAGCATCACGATGCTCGATGTTTGACGCTTCACAG |
| GGGCCTATCGTGGCCTCCCACGTCCTTCATCGGTTCCTGG |
| GGGCTTTCACAACCGACGTGACAAGCCGCCTACGAGCTCT |
| GGGGGCACCATCTCTGATGCTTTCCGGTGTATGTCAAGCC |
| GGTCTTGAGCGTGCTACTAAAACGCCCTATTCGGACTCGC |
| GGTCTTTCCGTCCTGCTGCGCGAAACGAGCATCTTTACTC |
| GGTTCGGTCCTCCACGACCTCTTACAGCCGCTTCAACCTG |
| GTAATCGGGTGACACCACGAGAGCGGAACAGTCGGAGGAATAA |
| GTACCGGAATATCAACCGGTTATCCATCGACTACGCCTGTC |
| GTACCTTTTATCCGTTGAGCGACGGCGCTTCCACAAGCCAC |
| GTACTTTTCACCATTCCCTCACGGTACTATCCGCTATCGGTCAC |
| GTAGGAGTCTGGGCCGTGTCTCAGTCCCAGTGTGGCCGGTCG |
| GTAGTTAGCCGGCGCTTCTTCTGCAGGTACCGTCACTTTC |
| GTATACATCGCCTTACGGCTTCGCACGGACCTGTGTTTTTA |
| GTATCGCAGCTCATTGTACCGGCCATTGTAGCACGTGTGCAGCCCAAGACATAA |

| |
|---|
| GTATTAGACCCCGTTTCCAGGGCTTGTCCCAGAGTGAAGGGCAGATT |
| GTCCTCTCGTACTAGGGACAGCCCTTCTCAAGACTCCTAC |
| GTCCTTCATCGGTTCCTGGTGCCAAGGCATCCACCGTGCGCCCTTAAAAACTT |
| GTCGATATGGACTCTTGGGGAAGATCAGCCTGTTATCCCCGGGGTACCTTTTAT |
| GTCGCCTTGGTGAGCCATTACCTCACCAACAAGCTGATAG |
| GTCGGAACTTACCCGACAAGGAATTTCGCTACCTTAGGATGGTTATAGT |
| GTCTCACACGTATACGGTTTGGCCTCATCCGGTTTCGCTC |
| GTCTCACAGTCAAGCTCCCTTGTGCACTTACACTCAACACCTGATTG |
| GTCTCCCGTGAGTCCCCAACACCCCGAAGGGCTTGCTGGCAAC |
| GTCTGCTTTCTGTGGCACTGTCCCGCGGGTCACCCCGGGTGGCCGTTAGCCATCAC |
| GTCTTGAGCGTGCTACTAAAAACGCCCTATTCGGACTCGC |
| GTGGGAGTCCTCATCTCGAAGCAGGCTTCCCGCTTAGATG |
| GTGTACAAGGCCCGGGAACGTATTCACCGCAGCAATGCTG |
| GTGTATGTCAAGCCTTGGTAAGGTTCTTCGCGTTGCGTCGAATTAAG |
| GTGTCTCTCAAACGAGCCGTTGACGTTTCGACTACGGGGGTCTTACCCTCTA |
| GTGTTACCGACTTTCGTGACGTGACGGGCGGTGTGTACAAG |
| GTGTTTTTAGTAAACAGTCGCTTCTCGCTGGTCTCTGCGG |
| GTTACTCTTTAGGAGGCAACCGCCCCAGTTAAACTACCCATCAGACACTG |
| GTTAGACATCCAGCACGACCAGACTGGTATTTCAACGACGACTC |
| GTTCCCTCCACACTGCCTATGTGTTCAGCAGCGGGTGACAG |
| GTTCGAAATGTAACCGGGCGTTTCCCCTACGCTATGACCACCGAAA |
| GTTCGGCGGCGTCCTACTCTCCCACAGGGTCCCCCCTGCA |
| GTTCGGCGGCGTCCTACTCTCCCACAGGGTCCCCCCTGCAGTACCATCGGCGCTGTA |
| GTTCTGTTCCGACGGGGCCTCGCGGTCCCGTCGGCGACGGCCATCCAT |
| GTTGTCTGTGCGACTCCACATCCTTTCCCACTTAGCGTACGCTTAG |
| GTTTACTGGCGCTTAAGTTCTCAGCTTCGCCACACCGAAATGTGACTAAC |
| GTTTCAGGTACTATTTCACTCCGCTCCCGCGGTACTTTTCACCATTC |
| GTTTCGCTCGCCACTACTCCCGGAATCACGGTTGTTTTCTCTTC |
| GTTTGATTGGCCTTTCACCCCTAACCACAGGTCATCCCCCAGGTTTTCAACC |
| GTTTGTATCGCTACTCATGCCTGCATTCTCACTCGTGAACCGTCCACAAC |
| GTTTTCAACCCTGGTGGGTTCGGTCCTCCACGACCTCTTAC |
| TAAGTTCTCAGCTTCGCCACACCGAAATGTGACTAACCGGTCCCCTTAACGTT |
| TACATCAACGGTTTCTGACTCGTCTCACGGCCGGCAGACCGTAAAAGAGAGAT |

| |
|---|
| TACCGACTTTCGTGACGTGACGGGCGGTGTGTACAAGGCCCGGGAACGTATTC |
| TACCTTTTATCCGTTGAGCGACGGCGCTTCCACAAGCCACCGCCGGATCACTAGTC |
| TACGGCTTCCCCACTCGGGTTAACCTCGCAACACACCGCAAACTCGCAGGCTCAT |
| TACTCATGCCTGCATTCTCACTCGTGAACCGTCCACAACTCGCTTCCGCGGCTGCTT |
| TAGGAGTCTGGGCCGTGTCTCAGTCCCAGTGTGGCCGGTCGCCCTCTCAGGCCGGCTA |
| TATACATCGCCTTACGGCTTCGCACGGACCTGTGTTTTTAGTAAACAGTCGCTTCT |
| TATCAACATATCTGGCGTTGACTTTTGGCACGCTGTTGAGTTCTCAAGGAA |
| TATCCAGTGCTCTACCTCCGGCAAGAAACACACGACGCTGCACCTAAATGCATTT |
| TATCCGCTATCGGTCACCAGGGAATATTTAGGCTTAGCGGGTGGTCCCGCCAGATT |
| TATCGACTGCAGACCCGGGGTTAAGCCCCGGGCTTTCACAACCGACGTGACAAG |
| TATGGTTGAGACAGTCGAGAAGTCGTTACGCCATTCGTGCAGGTCGGAACTTACC |
| TATTACGCACTCTTTCAAGGGTGGCTGCTTCTAAGCCAACCTCCTGGTTGTCTGT |
| TATTCACCGCAGCAATGCTGATCTGCGATTACTAGCGACTCCGACTTCATG |
| TATTCTCTACCTGACCACCTGAGTCGGTTTAGGGTACGGGCCGCCATGAAACT |
| TATTGCTGCAATGACACGACTTCGGCGGTACGCTTGAGCCCCGCTACATTGTCG |
| TCAAGCCTTGGTAAGGTTCTTCGCGTTGCGTCGAATTAAGCCACATGCTCCGCCGCTT |
| TCAATTCCTTTGAGTTTTAGCCTTGCGGCCGTACTCCCCAGGCGGGGCACTTAA |
| TCACGACACGAGCTGACGACAGCCATGCACCACCTGTACACCGACCACAAGGGGGGCACC |
| TCACGACGTTCTAAACCCAGCTCGCGTACCGCTTTAATGGGCGAACAGCCCAACC |
| TCATCCTTCACCGCCGCAGCTTTCGAACCTCGCAGATGCC |
| TCCACAACTCGCTTCCGCGGCTGCTTCACCCGGCACACGACGCTCCCCTACCCATCACA |
| TCCACACTGCCTATGTGTTCAGCAGCGGGTGACAGCCCATGACGACTGCCGGGTTT |
| TCCTTAGAAAGGAGGTGATCCAGCCGCACCTTCCGGTACGGCTACCTTGTTACGA |
| TCGCAGATGCCTGCAAGGGTCAGTATCCGGTATTAGACCCCGTTT |
| TCGGACTCGCTTTCGCTACGGCTTCCCCACTCGGGTTAAC |
| TCTCACACGTATACGGTTTGGCCTCATCCGGTTTCGCTCGCCACTACTCCCGGAAT |
| TCTTCAAAAGGCACGCAGTCACGAGACACCAAGCAAGCTT |
| TGCAGTCGGCTATGTGCTCCTTAGAAAGGAGGTGATCCAG |
| TGCATCAGGCTTTCGCCCATTGTGCAATATTCCCCACTGCTGCCTCCCGTAGGAGTCT |
| TGCGGATTTACCTACACACCGGCCTACACCCTTACCCCGGGACAACCACCG |
| TGCTGAAAGAGGTTTACAACCCGAAGGCCGTCATCCCTCACGCGGCGTCGCTGCATCA |
| TGTAACCGGGCGTTTCCCCTACGCTATGACCACCGAAACC |
| TGTGTTCAGCAGCGGGTGACAGCCCATGACGACTGCCGGGTTTCCCCATTCGGACACCC |

| |
|---|
| TGTGTTTTTAGTAAACAGTCGCTTCTCGCTGGTCTCTGCGGCCACCCCCAGCTCAGA |
| TGTTACTCACCCGTTCGCCACTAATCCCCACCGAAGTGGTTCATCGTTCGACTT |
| TTAAACTACCCATCAGACACTGTCCCTGATCCGGATCACGGACCCAGGTTAGACAT |
| TTAACCCAACATCTCACGACACGAGCTGACGACAGCCATGCACCACCTGTACACCGA |
| TTAGACATCCAGCACGACCAGACTGGTATTTCAACGACGACTCCACCCACACTGGCGTG |
| TTAGATGCTTTCAGCGGTTATCCCTCCCGAACGTAGCCAACCAGCCATGCCCTTGGCA |
| TTAGCATCACGATGCTCGATGTTTGACGCTTCACAGCGGGTACCGGAATATCAA |
| TTAGGATGGTTATAGTTACCACCGCCGTTTACTGGCGCTTAAGTTCTCAG |
| TTATCCATCGACTACGCCTGTCGGCCTCGCCTTAGGTCCCGACTTACCCTGGGCA |
| TTCAAAGGAACCTCAACCCGAACGGAGATCCGATCAGGTCGGGGTATCAACATAT |
| TTCACCGCCGGAGCTTTCGACCCTCGCAGATGCCTGCGAG |
| TTCCTTCTTGACCTTGCTCCGGGTGGGGTTTACCTAGCTGCCCAGGTCGCCCTG |
| TTCGACCCTCGCAGATGCCTGCGAGGGTCAGTATCCGGTATTAGACCCCGTTTCCAG |
| TTCGACTTGCATGTGTTAAGCACGCCGCCAGCGTTCGTCCTGAGCCAGGATCAAACTCT |
| TTCGGCGGCGTCCTACTCTCCCACAGGGTCCCCCTGCAGTACCATCGGCGCTGTAA |
| TTCGTCCCAATCGCCAGTCCCACCTTCGACAGCTCCCTCCCACAAGGGGTTGGG |
| TTCTCACTCGTGAACCGTCCACAACTCGCTTCCGCGGCTGCTTCACCCGGCACACGACG |
| TTCTTCGCGTTGCGTCGAATTAAGCCACATGCTCCGCCGCTTGTGCGGGCCCCCGTCAA |
| TTCTTCTGCAGGTACCGTCACTTTCGCTTCTTCCCTGCTGAAAGAGGTTTACAA |
| TTGTTTTCTCTTCCTGCGGGTACTGAGATGTTTCACTTCCCCGCGTTCCCTCCACA |
| TTTAATGGGCGAACAGCCCAACCCTTGGGACCGACTCCAGCCCCAGGATGCGACGA |
| TTTAGGGTACGGGCCGCCATGAAACTCGCTAGAGGCTTTTCTC |
| TTTCGCATGTCCTTCGCCTACATCAACGGTTTCTGACTCGTCTCACGGCCGGCAGA |
| TTTCTGACTCGTCTCACGGCCGGCAGACCGTAAAAGAGAGATCCCACAACCCCGCACA |
| TTTTAGCCTTGCGGCCGTACTCCCCAGGCGGGGCACTTAATGCGTTAGCTGCGGCAC |
| TTTTCTCGACAGCATAGGATCATCCACTTCACCACAATCGGCTCGGCATCAGGTCT |
| TTTTGAGATTCGCTCCACCTTGCGGTATCGCAGCTCATTGTAC |
| TTTTGAGATTCGCTCCACCTTGCGGTATCGCAGCTCATTGTACCGGCCATTGTAGCACGT |
| TTTTTCGGCCTGTGGTTTACGAGATCATGGCCGCTTCCTC |
